# Transcriptional induction and mechanical propagation of a morphogenetic wave

**DOI:** 10.1101/430512

**Authors:** Anais Bailles, Claudio Collinet, Jean-Marc Philippe, Pierre-François Lenne, Edwin Munro, Thomas Lecuit

## Abstract

Tissue morphogenesis emerges from coordinated cell shape changes driven by actomyosin contractions. Patterns of gene expression regionalize and polarize cell behaviours by controlling actomyosin contractility. Yet how mechanical feedbacks affect tissue morphogenesis is unclear. We report two modes of control over Rho1 and MyosinII activation in the *Drosophila* endoderm. First, Rho1/MyoII are induced in a primordium via localized transcription of the GPCR ligand Fog. Second, a tissue-scale wave of Rho1/MyoII activation and cell invagination progresses anteriorly. The wave does not require sustained gene transcription, and is not governed by regulated Fog delivery. Instead, MyoII inhibition blocked acute Rho1 activation and propagation, revealing a mechanical feedback driven by MyoII. Last, we identify a cycle of 3D cell deformations whereby MyoII activation and invagination in each row of cells drives adhesion to the vitelline membrane, apical spreading, MyoII activation and invagination in the next row. Thus endoderm morphogenesis emerges from local transcriptional initiation and a mechanically driven wave of cell deformation.

---

The understanding of how embryonic forms arise during development requires accounting how controlled mechanical operations emerge from pre-established information and dynamic molecular interactions. The current mechanistic framework explains how genetic and biochemical information controls cellular mechanics^1,2^. Developmental information regionalizes and polarizes cellular behaviours. The early *Drosophila* embryo, where the expression of patterning genes defines cell behaviour, illustrates this. Here the expression of the mesoderm transcription factors Twist and Snail induces apical cell constriction and tissue invagination in a ventral region of the embryo^3^.

A large body of work highlighted how these cellular behaviours arise from spatial and temporal control over actomyosin contractility^1,4^. For instance, activation of non-muscle Myosin II (MyoII) in the medio-apical region of cells drives their apical constriction and tissue invagination^5–8^. The small GTPase Rho1 activates MyoII through the downstream kinase Rok^9–12^ and is itself controlled by different signalling pathways, including G protein coupled receptors (GPCRs)^13,14^. During gastrulation in *Drosophila* the localized transcription and secretion of the GPCR ligand Fog controls Rho1 activation and MyoII-dependent apical constriction^15,16^. Thus developmental patterning controls actomyosin contractility and tissue dynamics through control of Rho signalling. Yet, during morphogenesis, actomyosin networks and Rho GTPases display remarkable dynamics that are not strictly governed by upstream genetic programs. Actomyosin networks are pulsatile in a wide variety of species^5,6,17 7,12,18–20^, they display remarkable flow dynamics^17,18,21,22^ while cortical pulses and waves of RhoA activity emerge at the cortex of large cells^12,23,24^. Furthermore actomyosin networks can sense and respond to the mechanical environment^25–28^ which itself can provide both spatial and temporal information. How genetically encoded information and mechanical control integrate to coordinate actomyosin contractility and tissue level morphogenesis is still largely unexplored. We address this in the early *Drosophila* embryo and report a mechanically driven wave of MyoII activation and cell deformation during morphogenesis of the posterior endoderm.

## A wave of MyoII activation and cell invagination travels anteriorly

The *Drosophila* posterior endoderm consists of a circular domain of cells at the posterior pole of the embryo that undergoes invagination and movement towards the anterior, driving germ-band extension (Fig.1a)^29,30^. MyoII-dependent apical constriction drives local endoderm invagination^15,31,32^, but how the endoderm moves towards the anterior remains unknown. To investigate this we imaged MyoII (using an mCherry-tagged MyoII Regulatory Light Chain, MRLC::mCherry) together with a marker of cell contours (E-cadherin::GFP) on the dorsal posterior side of embryos (Fig.1b and Movie 1). During the first 6 minutes MyoII is activated in the medial-apical region of cells within a spatially defined region (the endoderm primordium) comprising 7±1 rows of cells from the posterior pole (Fig.1b top and 1d). Surprisingly, in the next 25 minutes, apical MyoII activation propagates anteriorly across a domain of 8±1 rows of cells hereafter called the propagation region (Fig.1b middle-bottom and 1d). Upon MyoII activation, cells incorporate in the invaginating furrow and constrict (Fig.1c, Ext.Fig.1g-i), thereby leading to forward expansion of the invagination. Thus, the region where MyoII activation and cell invagination occur is dynamic. This contrasts with the *Drosophila* mesoderm, where apical constriction occurs in a fixed domain defined by the expression of *twist* and *snail*^2^.

**Figure 1.**
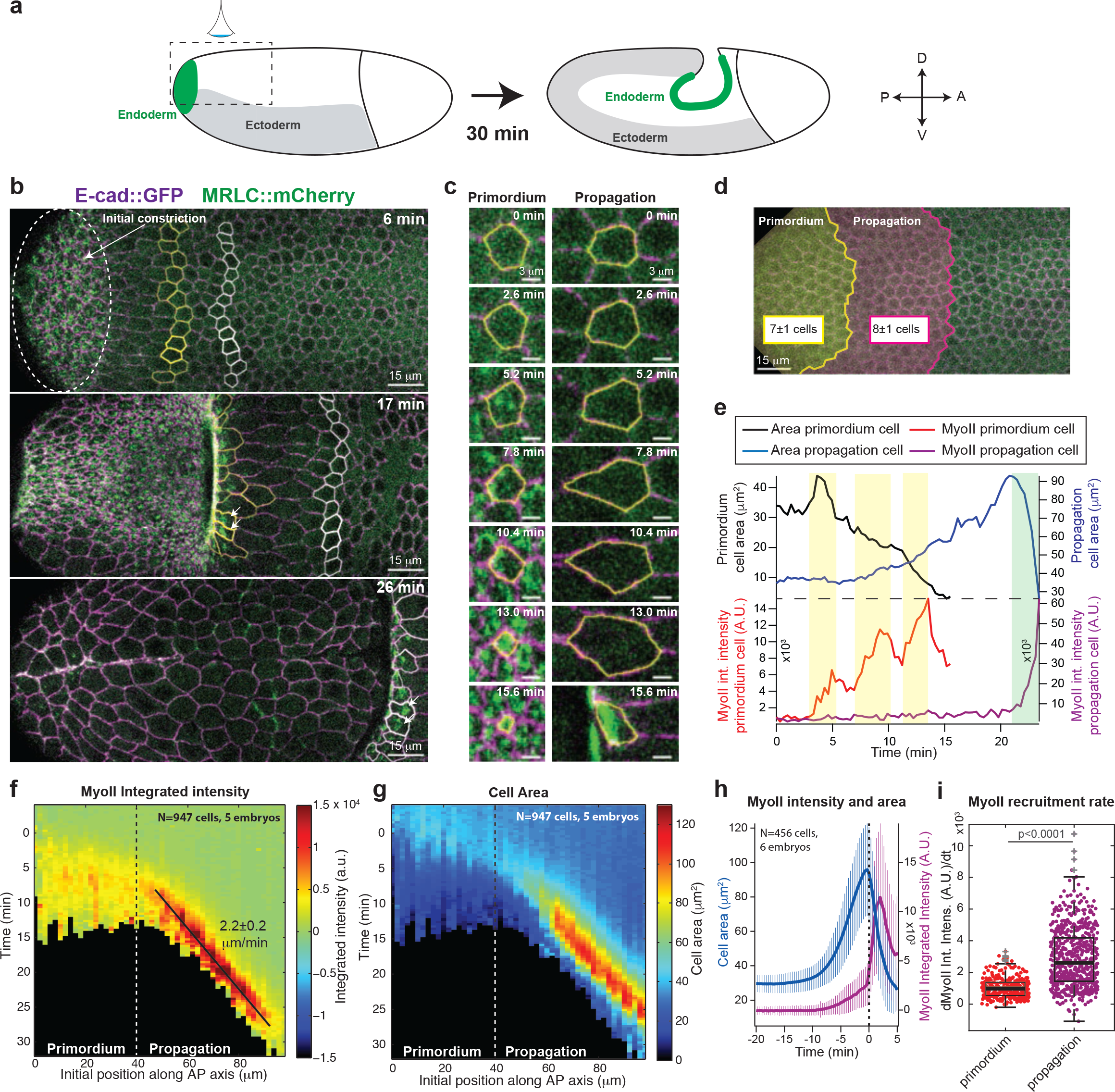
Propagation of MyoII activation during morphogenesis of the posterior endoderm. (**a**) Cartoon of endoderm morphogenesis during embryonic axis extension. On the left: the ventrolateral ectoderm (in grey) and the posterior endoderm (in green) are depicted at the onset of axis extension and 30’ later. The dotted box indicates the region of imaging. (**b**) Time-lapse confocal images of endoderm morphogenesis. The dashed oval marks the primordium region and yellow and white contours mark cells of the propagation region where MyoII is activated at 17min and at 26min, respectively (indicated by arrows). (**c**) Close up views of a cell in the primordium (left) and propagation (right) regions. (**d**) The primordium and the propagation region mapped on the dorsal epithelium at the onset of gastrulation. (**e**) Measurements of projected apical cell area and MyoII integrated intensity of the cells in **c**. The yellow and green boxes highligth steps of MyoII recruitment in the primordium cell and the cell in the propagation region respectively. (**f-g**) Kymograph heat-maps of MyoII integrated intensity (**f**) and projected apical cell area (**g**) obtained from cell tracking in the endoderm. The dashed line indicates the primordium and propagation regions and the black solid line indicates the constant travelling speed of the MyoII wave. N=947 cells from 5 embryos. (**h**) Average time course of the projected apical cell area and MyoII integrated intensity in cells in the propagation zone (N=456 cells from 6 embryos). Time 0 is defined for each cell when MyoII intensity reaches a threshold (see methods). (**i**) Measurements of MyoII recruitment rate in cells of the primordium and propagation (N=288 cells for the primordium and 456 cells for the propagation region from 6 embryos). In **b**, **e**, **f**, **g**, time 0 is the onset of endoderm morphogenesis (see methods). Mean ± SD are shown in **h**.

We tracked cells over time and plotted measured MyoII intensity and apical cell shape as a function of time and initial cell position along the AP-axis (Fig.1f-g and Ext.Fig.1e-f, see Methods and Ext.Fig.1a-d). Heat-map kymographs show that the MyoII activation wave travels with a constant speed of 2.2±0.2 μm/min (corresponding to one cell every ~3 min, Fig.1f) and is preceded by a wave of anisotropic cell deformation where cell apical areas (measured as their projection on the imaging plane, Ext.Fig.1g) increase along the AP-axis before invagination (Fig.1g,h and Ext.Fig.1e-f). Strikingly, while cells in the primordium recruit MyoII and constrict apically in a stepwise manner (Fig.1c,e and Movie 2), cells in the propagation region recruit MyoII with kinetics significantly faster than the primordium (Fig.1,c,e,i). This suggests different mechanisms of MyoII activation in the primordium and in the propagating regions.

We further imaged Rho1 activation with a Rho1-GTP biosensor^10^, and a Rok::GFP^9^ together with MyoII::mCherry and found that both Rho1-GTP and Rok are activated together with MyoII in the propagation domain (Ext.Fig.2 and Movies 3-4). Thus a wave of Rho1, Rok and MyoII activation propagates across the dorsal epithelium. This led us to investigate the underlying mechanisms.

## The MyoII wave does not depend on propagation of gene expression

The medio-apical activation of MyoII in the endoderm primordium depends on localized transcription and secretion of Fog^15,16^, which is controlled by terminal patterning through the receptor tyrosine kinase Torso (Tor) and its target transcription factors Huckebein (Hkb) and Tailless (Tll)^15,33,34^ (Ext.Fig.3a). In mutants for *tor* or *fog* or for *concertina*, the G-protein Gα_12/13_ downstream of GPCRs^16^, MyoII apical activation was abolished in the primordium, as expected, but also in the propagation region (Ext.Fig.3b and Movie 5). We thus tested whether the expression of *hkb* and *tll* and their target *fog* propagates anteriorly within the dorsal epithelium to activate MyoII in cells of the propagation region (Fig.2). Mid-sagittal sections of immunolabelled embryos revealed that Fog is present in the primordium where MyoII is activated (Fig.2a, primordium contraction stages). The *fog* and *tll* expression domains did not expand over time and their anterior boundaries were in increasingly deeper positions of the invagination as new cells incorporated (Fig.2a and Ext.Fig.3c, MyoII propagation stages). In contrast MyoII was enriched at cell apices at the invaginating front at all stages (Fig.2a), reflecting propagation of its activation.

**Figure 2.**
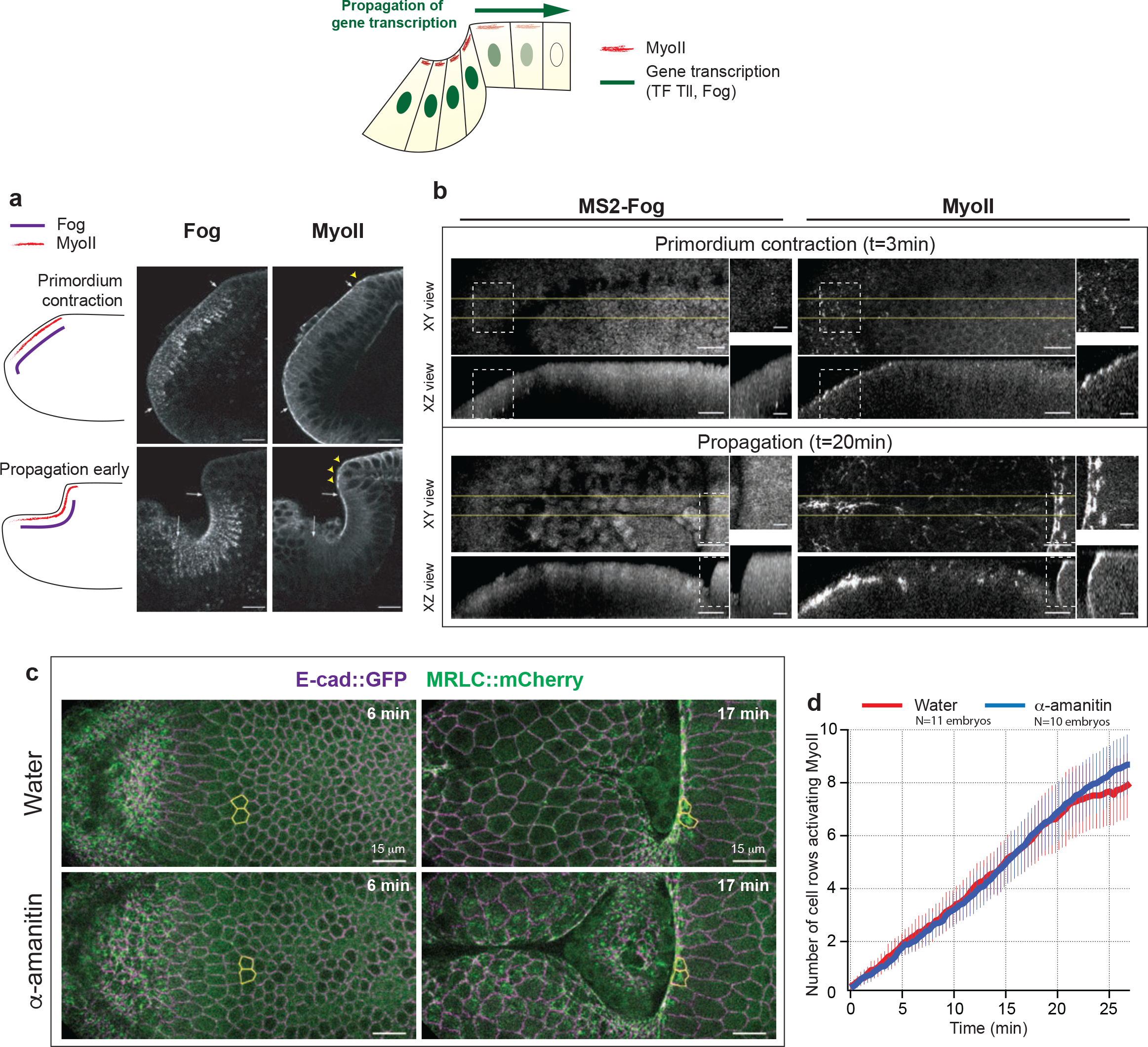
MyoII activation in the primordium but not in the propagation region is controlled by gene transcription. Top: Cartoon of the dorsal epithelium in a sagittal section describing the hypothesis of spatio-temporal changes in gene transcription controlling MyoII propagation. (**a**) Sagittal sections of embryos stained for Fog and MyoII at primordium contration and wave propagtion stages. On the left a cartoon representation of the distribution of the indicated molecules. (**b**) *fog* expression in the posterior endoderm visualized with the MS2-MCP system^35^ and MyoII recruitment in living embryos. Top and side views are shown for the indicated time points. White dashed boxes indicate the positions of the close-ups on the right and the yellow lines the region where side-views were generated. *fog* is expressed in the primordium but not in the propagation zone. (**c**) Posterior endoderm morphogenesis in control (water) and α-amanitin injected embryos. Yellow contours mark cells in the propagation region. (**d**) Quantifications of the MyoII activation wave in water and α-amanitin injected embryos. The number of cell rows aligned along the ML-axis activating MyoII in the propagation region over time was measured. N=11 for water and N=10 for a-amanitin injected embryos. Mean ± SD are shown.

We also monitored *fog* transcription in living embryos together with MyoII using the MS2/MCP system^35^. Fog transcripts containing MS2 stem loops in the 3’UTR (see Methods) appeared as bright GFP dots in the nuclei marked with His2B::RFP (Ext.Fig.3d). *fog* was expressed in the primordium at the beginning of gastrulation but was strikingly absent from the propagation zone (Fig.2b and Movie 6). We concluded that while MyoII activation in the primordium depends on localized transcription of the GPCR ligand Fog, the subsequent wave is not associated with a wave of *fog* transcription in the propagation zone. Transcription of other genes might underlie a relay mechanism to propagate MyoII activity. To test this, we injected α-amanitin, a potent inhibitor of RNA Polymerase-II. Injection at the end of cellularization led to a rapid disappearance (within 2-3 mins) of nuclear foci of nascent Fog transcripts (Movie 7) while the wave of MyoII proceeded at normal velocity (Fig.2c,d and Movie 8). Two processes requiring acute transcription were affected in these conditions: 1) cell divisions in mitotic domains (dependent on *cdc25^string^* transcription^36^) were blocked and 2) cell intercalation requiring *even-skipped* and *toll2,6,8* transcription^37^ in the ventro-lateral ectoderm were inhibited (Ext.Fig.3e and Movie 8). These experiments demonstrate that while MyoII wave is initiated by *fog* transcription in the primordium, its progression depends on a relay mechanism independent of transcription.

## Delivery of Fog produced in the primordium does not govern the MyoII wave dynamics

Fog is thought to act through paracrine signalling^15^. Thus, in principle, the spread of Fog away from the primordium, where it is produced and secreted, could underlie propagation of Rho1 and MyoII activity. Consistent with this, immuno-stained Fog protein was detected, albeit at low levels, at the cell apices in the propagation zone close to the furrow (Ext.Fig.4a,b). Simple models for diffusive spread of secreted ligands predict a wave front that flattens away from the source, which is inconsistent with the kinematics observed experimentally (Fig.1f). Building upon earlier work^38,39^, a model that couples Fog diffusion and saturable receptor binding to MyoII activation can produce a travelling wave of sharp MyoII activity (Fig.3a,b and Supplementary Information). This model predicts that increasing the rate of ligand production increases the speed of wave propagation (Fig.3b). We thus experimentally increased Fog expression in the primordium using 2 extra copies of *fog* under the control of the *hkb* promoter^40^, and measured MyoII activation (Fig.3c, Ext.Fig.4c-e and Movie 9). This resulted in earlier recruitment of MyoII in the primordium (Ext.Fig.4c), consistent with increased Fog dose. MyoII levels were increased in the primordium and to a lesser extent in the propagation zone (Ext.Fig.4e). Contrary to model predictions, despite increased Fog levels, the rapid activation of MyoII in the propagation region occurred at the same time at similar distances from the primordium as in controls (Ext.Fig.4d), such that no change in wave speed was observed (Fig.3d). This suggested that source-limited diffusive spread of Fog is not the rate-limiting factor for MyoII propagation.

**Figure 3.**
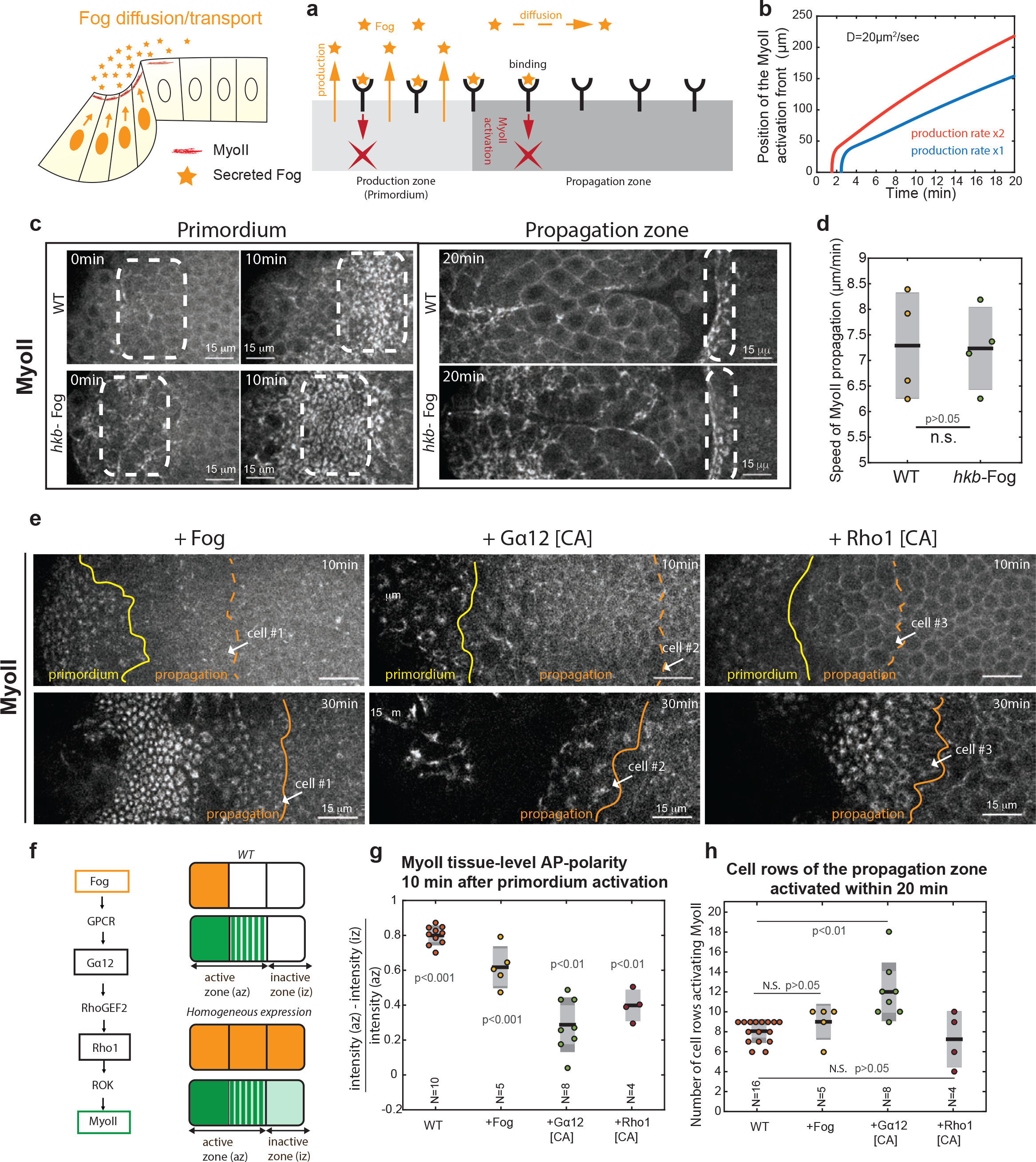
Fog diffusion and/or patterned Fog signaling do not control MyoII wave propagation. Top left: Cartoon describing the hypothesis of secreted Fog diffusion/transport from the primordium controlling MyoII propagation. (**a**) Schematic representation of the model of Fog secretion, diffusion, receptor binding and MyoII activation (see details in Supplementary Information). The red crosses represent activated MyoII. (**b**) Position of MyoII activation front over time obtained from simulations using the model described in **a** for the indicated diffusion constant (D), a given production rate (blue) and its double (red). (**c**) Time-lapse of MyoII in the primordium and propagation region of WT and embryos overexpressing Fog in the primordium (hkb-*fog*). The dashed line boxes mark the primordium and the activated cells in the propagation region respectively. (**d**) Speed of MyoII propagation in the reference frame of the microscope (measured by a linear fit covering the propagation phase (10-30 min from the onset of endoderm morphogenesis), see methods). Individual data points are superimposed to box plots (dark line: mean, light grey zone: 95% s.e.m., darker grey zone: s.d.). N=4 embryos each. N.S. indicates a P>0.05 from a Mann-Whitney test. (**e**) MyoII pattern in an embryo where Fog (left), a constitutively active form of Gα12/13 (middle) or a constitutively active form of Rho1 (right) are expressed uniformly. The yellow and orange lines denote the limits of the primordium and propagation regions respectively, defined by manual tracking. The white arrows point the same cell in the two time points. (**f**) Left: Pathway of MyoII activation by Fog in endoderm cells. Right: Schematic of WT and uniform expression of Fog in the embryo. Fog expression pattern is shown in orange and the resulting MyoII pattern in green. The MyoII active and inactive zones used to calculate the MyoII tissue level polarity in **g** are indicated. (**g-h**) Quantifications of MyoII tissue-level polarity (**g**) and of the number of cell rows along the ML-axis activating MyoII in the propagation region in embryos of the indicated conditions. (**h**) Individual data points are superimposed to box plots as for **d**. N indicates the number of embryos. For **g**, P<0.01 and P<0.001 for each condition are the p-values of a one-sample Student test against the null hypothesis that the data come from a normal distribution with mean equal to 0 (i.e. that the tissue has no tissue-level polarity of MyoII). For **h**, P>0.05 and P<0.01 from a Mann-Whitney test.

A wave of Fog signalling patterned by other mechanisms (e.g. ligand transport or GPCR regulation) might activate MyoII in cells of the propagation region via GPCRs/Rho1 signalling. To test this we overexpressed Fog uniformly in the embryo, overriding the endogenous Fog pattern (Fig.3e,f, Movie 10 and Ext.Fig.4f). Under these conditions, although MyoII levels in the dorsal epithelium were slightly increased, the tissue-level pattern, with high MyoII in the posterior endoderm and low MyoII more anteriorly, was preserved (Fig.3g). This pattern was not due to endogenous *fog* since uniform Fog expression in *fog−/−* embryos yielded similar results (Ext.Fig.4g,h). Importantly, high MyoII levels propagated at similar speeds and across a similar number of cell rows as in WT embryos (Fig.3e,h). Bypassing GPCR regulation also failed to prevent wave propagation. Uniform expression of the constitutively active Gα_12/13_ and Rho1 led to some (although lower) MyoII tissue-level polarity and anterior propagation of high MyoII levels still occurred (Fig.3e middle and right, 3g,h and Movie 10). Expression of Gα_12/13_[CA] led to propagation across more cell rows possibly as a result of the extensive tissue folding associated with this perturbation. Altogether, these data argue that patterned Fog signalling does not determine the dynamics of the MyoII wave. Fog signalling may be required but MyoII activation is not governed solely by Fog-dependent Rho1 activation in the propagation zone.

## Mechanical regulation of the Rho1/MyoII activation wave

Since MyoII can respond to mechanical stimuli^25,27^, we considered the possibility that mechanical stress (either acting directly on MyoII concentration/stabilization^41^ or indirectly through mechanosensitive signalling^42,43^) could be involved in propagating MyoII activation across the tissue. Consistent with this, before recruiting MyoII, cells in the propagation region are deformed anisotropically (Fig.1c, 1g and Ext.Fig.1e,f), and subjected to anisotropic stress, as indicated by laser ablations (Ext.Fig.5). To test whether MyoII-dependent tissue deformation is required to propagate the Rho1-GTP wave, we injected embryos with a Rok inhibitor (H-1152) before gastrulation and monitored the dynamics of Rho1-GTP. Following injection, Rho1 activation within the primordium (where Fog is expressed and secreted) was unaffected, but the propagation of Rho1 activation was sharply reduced to ~4 cell rows during the next 25-30 min (Fig.4a,b and Movies 11-12). This indicates that MyoII activity in the primordium is required for normal propagation of Rho1-GTP, and hence MyoII activation.

**Figure 4.**
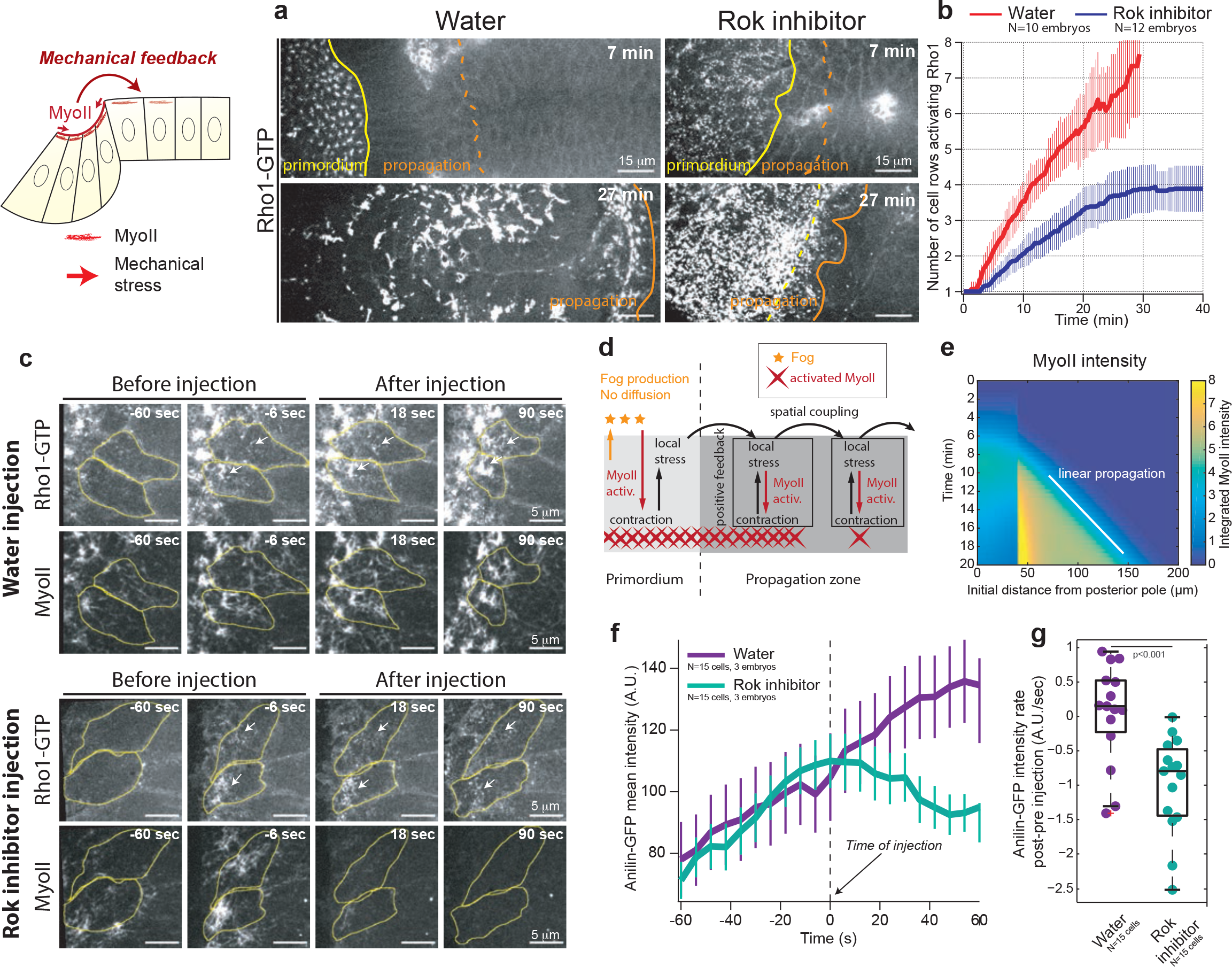
The wave of MyoII is mechanically regulated and a feedback from MyoII promotes Rho1 activation in cells. Top left: Cartoon describing the hypothesis of a mechanical feedback controlling MyoII propagation. (**a**) Time-lapse stills of Anillin(RBD) (Rho1-GTP sensor) in water and H-1152 (Rok inhibitor) injected embryos. Embryos were injected at end of cellularization. The yellow and orange lines denote the limits of the primordium and propagation regions, respectively. (**b**) Quantifications of the wave of Rho1 activation as in Fig. 2d. N=10 for water and N=12 for H-1152 injected embryos. Mean ± SD between different embryos are shown. (**c**) Stills from high resolution time lapses of Rho1 sensor (Anillin(RBD)::GFP) and MyoII in water (top) and H-1152 (Rok inhibitor) injected embryos (bottom) during propagation. Cell contours are outlined in yellow and the white arrows mark accumulations of Rho1-GTP. (**d**) Schematic representation of the hypotheses underlying the model used to study the effects a stress-based feedback on MyoII (see Supplementary Information). The red crosses represent activated MyoII. Fog is produced in the primordium where it activates MyoII but it is not allowed to diffuse. Instead, stress locally activates MyoII and propagates within the tissue. (**e**) Kymograph heat-map of activated MyoII quantity (the concentration of MyoII integrated over the unit volume taking into account its local deformation, see Supplementary Information) from simulations of the model in **d**. (**f-g**) Quantifications of the mean intensity of Rho1 sensor before and after injections. In **f** average mean intensity measurements over time are shown and in **g** quantifications of the Anillin(RBD)::GFP recruitment rate difference (post-pre injection) following Rok inhibitor injection. N=15 cells from 3 embryos in both water and H-1152 injections. P<0.001 from a Mann-Whitney test. Mean ± SD different cells from multiple embryos are shown in f and box plots overlaid to individual data in **g**. In **a** time 0 is the onset of endoderm morphogenesis, in **c** and **f** time 0 is the time of injection.

Next, we used two approaches to perturb mechanically the dorsal epithelium (but not the genetic identity of endodermal cells) and looked at the effects on MyoII propagation. Using laser-mediated tissue cauterizations^29,44^ we generated mechanical fences whereby cells are stitched to the fixed vitelline membrane surrounding the embryo. Medio-lateral dorsal cauterizations at ~30% egg length, ~20-30 cells anterior to the primordium, introduced mechanical resistance to the anterior movement of the posterior endoderm (Ext.Fig.6a middle-bottom and Movie 13). In dorsalized embryos, laid by *dorsal* mutant mothers, DV polarity is abolished and MyoII-dependent apical constriction in the endoderm primordium is rotationally symmetric, blocking anterior movement of the posterior endoderm (Ext.Fig.6a right-bottom and Movie 14). In both cases the MyoII activation wave slowed down (Ext.Fig.6b and Movie 13) and cells recruited higher levels of MyoII compared to WT (Ext.Fig.6c). The rate of cell apical constriction immediately following MyoII recruitment in the propagation region was significantly reduced despite higher MyoII levels (Ext.Fig.6d), indicating a higher resistance to invagination. Importantly, similar perturbations by mechanical fences in the lateral ectoderm did not affect subcellular MyoII activation^29^ revealing a unique mechanical sensitivity of posterior endodermal cells. Altogether, these experiments reveal specific mechanical control over MyoII levels and wave speed in the propagation region.

## MyoII-based positive feedback and spatial coupling are required for the activation wave

We hypothesized that wave propagation depends on a mechanical relay whereby cells ahead of the furrow activate MyoII in response to either tensile or shear stress and in turn invaginate, thereby propagating to anterior cells global tissue deformation, stress and MyoII activation. We used a 1D toy model to abstract and highlight minimal ingredients required to generate a wave of MyoII activity at constant speed through dynamic coupling of mechanical force and chemical activation. We considered that local contractile stresses produced by MyoII are transmitted by stretching the apical cortex against a frictional resistance and that the transmitted stress activates more MyoII^45^ (Fig.4d and Supplementary Information). For simplicity and generality, we modelled a direct dependence of MyoII activation on stress, while remaining agnostic about the underlying details or the exact nature (tensile or shear) of the stress. Simulations (Fig.4e) confirmed that two key elements are sufficient for propagation of MyoII activation at constant speed: 1) a strong non-linear positive feedback, in which MyoII locally generates stress and stress locally activates MyoII, giving rise to bistable dynamics; 2) a spatial coupling mechanism tuned by the mechanical properties of the system, in which deformation in one cell generates stress within finite distance, thereby triggering MyoII activation further away.

We tested whether these two elements are involved during the propagation of Rho1-GTP/MyoII activity. We assessed whether contractility is required to sustain Rho1 activation in cells following injection of the Rok inhibitor H-1152 during propagation. This immediately blocked the propagation of Rho1 activation across the tissue (Ext.Fig.7 and Movie 15) and the rapid increase of Rho1 activity in cells in the process of activation (Fig.4c, 4f-g and Movie 16). This confirmed the existence of a local positive feedback loop of contractility acting directly or indirectly on Rho1 activation. These results reveal the existence of both a cell-intrinsic (local amplification) and a cell-extrinsic (due to spatial coupling) feedback of contractility onto Rho1GTP.

## A mechanically relayed cycle of 3D cell deformations underlies propagation of the MyoII wave

To investigate the mechanical basis for spatial coupling, we imaged embryos in sagittal cross-section and identified a characteristic sequence of 3D cell deformations that accompanies the MyoII wave (Fig.5a,b and Movie 17). Cells anterior to the furrow were initially tall with small apices (Fig.5a magenta cell in insets). As the advancing furrow reached more anterior cells, their basal ends became narrower and moved anterior and upward, while their apical surfaces spread onto the vitelline membrane along the AP axis (Fig.5a yellow cell in insets). The apices then constricted again when cells joined the furrow (Fig.5a and Movie 17). MyoII activation was initiated in cells as they spread their apical surfaces (Ext.Fig.8 and Movie 18) and increased as cells detached from the vitelline membrane. Imaging of adherens junctions further documented the spreading of apical surfaces against the vitelline membrane (Fig.5c and Movie 19). Adherens junctions moved apically, first at the posterior, then at the anterior of each cell. As they reached the vitelline membrane, their anterior velocity slowed down before immobilization (Fig.5d,e and Movie 19). This sequence of events argues that cells are pushed towards the anterior and upwards against the vitelline membrane, and that subsequent frictional coupling of cell apices to the vitelline membrane resists anterior translation of the cell apex. We injected fluorescent Dextran in the perivitelline space to assess the free space between the apical surface and the vitelline membrane. Strikingly, Dextran was excluded ahead of the furrow (Fig.5f, Ext.Fig.9a,b and Movie 20), indicating that cells were indeed pushed against the vitelline membrane.

**Figure 5.**
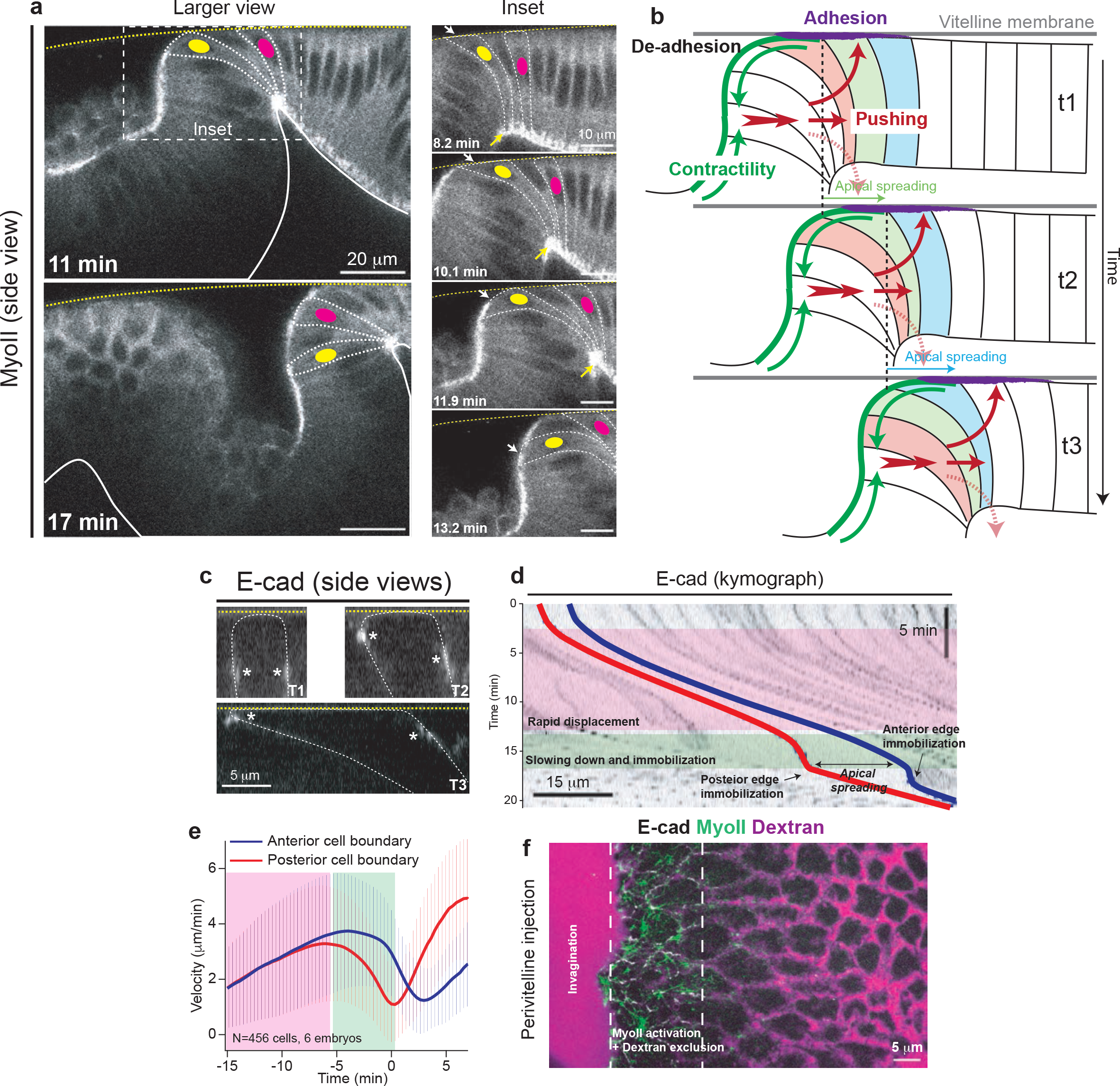
A 3D cycle of cell deformations and cell adhesion to the vitelline membrane accompanies the wave of MyoII and sustains MyoII activation. (**a**) Side views of MyoII recruitment and cell deformations in the propagation zone during posterior endoderm morphogenesis. Larger views are shown on the left and closeup of the white dashed boxed region are shown on the right. The dashed white lines highlight cell contours, the yellow dashed line the vitelline membrane (identified from its autofluorescence) and the solid white line the basal side of the blastoderm epithelium. The yellow and magenta dots mark 2 cells of the propagation zone. The white and the yellow arrows in the close up views indicate the apical and basal side of cells respectively. (**b**) Proposed model of the 3D mechanical cycle associated to the MyoII wave. At time tl MyoII contractility (in green) is induced in the posterior invagination and in the cell at boundary of the furrow (red cell) where it is anchored to the vitelline membrane by apical adhesion (labeled in purple). This leads to cell detachment and invagination and resulting pushing forces induce apical cortex spreading and adhesion to the vitelline membrane in the next row of cells (green cell at time t2). As a result the adhesion region is displaced anteriorly. At time t2 MyoII is induced in the green cell at the boundary of the furrow re-iterating the cycle to the next cell (blue cell at time t3). The red arrows represent pushing forces; the solid arrows contribute to deformations in the cycle while the dashed arrows do not. (**c**) Representative side views of E-cad junctions during the process of apical spreading of a cell in the propagation zone. The yellow dashed line labels the vitelline membrane (identified from its autofluorescence) and the white dashed line the cell outline. The white asterisks label the cell junctions. (**d**) A representative kymograph illustrating the movement (relative to the vitelline membrane and the deformation along the AP-axis of a cell in the propagation zone. E-cad is used to label cell contours. The red and blue traces label the posterior and anterior edges of the cell respectively. The pink and green boxes highlight the phases of rapid cell displacement and immobilization of the posterior edge. (**e**) Average velocity (relative to the vitelline membrane) of the posterior and anterior edges of cells in the propagation zone. Time 0 is defined for each cell as the time of max projected area increase rate. Mean ± SD between different cells are shown. N=456 cells from 6 embryos. (**f**) Still image from time laspe of dextran injection in the perivitelline space during MyoII propagation. The white dashed lines indicate the region of MyoII activation and dextran exclusion.

These observations led us to propose a 3D morphogenetic cycle as follows (Fig.5b). First, cells just anterior to the furrow initiate activation of MyoII while in contact with the vitelline membrane. MyoII activation persists as these cells detach from the vitelline membrane and invaginate and gives rise to tissue-scale contractile forces (Fig.5b, green arrows). These contractile forces result in normal pushing forces towards the anterior (Fig.5b, thick red arrow) due to cytoplasm incompressibility as proposed for the mesoderm^46^. As a result, the more basal regions of cells anterior to the furrow are displaced towards the anterior and compressed laterally (as revealed by a decrease in width in lateral cross-sections), displacing their basal cytoplasm upwards (Fig.5a). This in turn induces cells to push against and spread their apical surfaces onto the vitelline membrane, eventually activating MyoII to complete the cycle.

Remarkably, we observed a tight correlation between the speeds of 3D cell deformation and MyoII activation waves observed in WT, *dorsal* mutants and cauterized embryos, supporting a model in which MyoII both drives and is in turn induced by the cycle of cell deformation and attachment to the vitelline membrane (Ext.Fig.9c-e). Thus blocking MyoII activation should arrest all steps of this travelling cycle. This is indeed what we observed when we inhibited Rok during MyoII propagation (Ext.Fig.9f and Movie 21). Therefore the 3D travelling cycle requires sustained MyoII activity.

## Wave propagation requires integrin dependent adhesion to the vitelline membrane

A key feature of this model is the local resistance of apical movement against the vitelline membrane, leading to apical spreading before MyoII activation in cells anterior to the furrow. In occasional cases, cells appeared to resist detachment from the vitelline membrane before being incorporated in the invagination (Ext.Fig.9g and Movie 22), suggesting that this local resistance is mediated by specific adhesion to the vitelline membrane. Interestingly, αPS3 integrin (*scab* in *Drosophila*) is expressed in the propagation region and is required for germband extension^47,48^. RNAi against *scab* resulted in a striking change in cell velocities in the propagation zone (Fig.6a,b and Movie 23), consistent with a role of αPS3/Scab in cell adhesion to the vitelline membrane. In WT embryos cells just anterior to the furrow moved anteriorly with slow velocities. In *scab* RNAi embryos however, cells moved faster towards the anterior compared to WT, and meanwhile tissue invagination was delayed and shallow. This pattern of movements suggests that apical cell surface adhesion to the vitelline membrane resists pushing forces towards the anterior and thereby facilitates tissue invagination. We also noticed that, in a subsequent phase, once an invagination had formed in *scab* RNAi embryos, cells moved towards the posterior into the invagination. This suggests that adhesion may provide dynamic apical-anterior anchorage to the MyoII-dependent contractile forces promoting the sustained anterior displacement of the furrow.

**Fig. 6.**
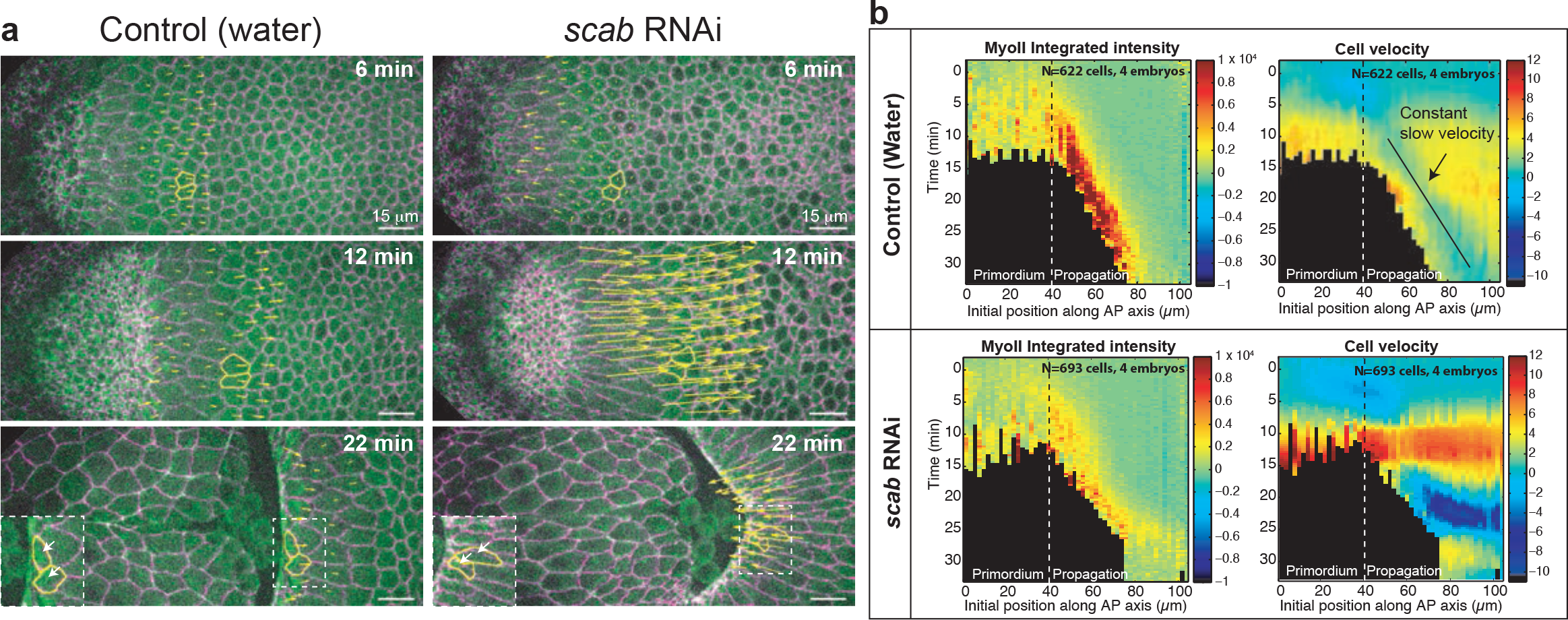
Integrin-dependent adhesion underlies posterior endoderm movement and MyoII activation during wave propagation. (**a**) Time-lapse stills of a control (water injected) and an embryo injected with dsRNA against *scab*. MyoII::mCherry and cell contours labeled with Ecad::GFP are shown. The yellow arrows are proportional to the velocities of cells in the propagation region. Two representative cells are marked in yellow. (**b**) Kymograph heat-maps of MyoII integrated intensity (left) and cell velocities (right) in the indicated conditions. The dashed line indicates the boundary between the primordium and the propagation region. N=622 cells from 4 water and 693 cells from 4 *scab* RNAi injected embryos.

Importantly, MyoII levels in cells of the propagation zone were significantly reduced during wave progression in *scab* RNAi embryos (Fig.6a,b), showing a role for integrin-based adhesion in MyoII activation in these cells.

In summary, local forces produced by MyoII drive an invagination that pushes cells towards the anterior, and their subsequent adhesion to the vitelline membrane generates forces that lead to more anterior activation of MyoII and cell invagination. This cycle of events is iteratively repeated linking MyoII contractility in the invaginating furrow to the anterior-wards movement of the endoderm (Fig.5b).

## Discussion

We describe a tissue-level wave of MyoII activation and cell invagination underlying the morphogenesis of the posterior endoderm during *Drosophila* germband extension. This morphogenetic wave is initiated by a spatially defined genetic input, in which a Fog signal transcribed within the endoderm, activates Rho1 and MyoII-dependent apical constriction and tissue invagination. The wave subsequently propagates via a mechanochemical relay in which local invagination drives cell displacement, adhesion, stress-induced contraction and further invagination/displacement. Stress-dependent activation of Rho1/MyoII provides both feedback to amplify local contractility, and spatial coupling to transmit contractile activity, leading to a traveling wave of contractility akin to trigger waves in excitable media^49,50^. Mechanical stress could activate MyoII in multiple ways: either directly via e.g. stress-dependent MyoII motor stabilization within the actomyosin cortex^41^ or more indirectly, via sensing of membrane tension^51^, and/or mechanosignalling^42,43^. In particular we uncover a key role of Integrins in wave propagation. Integrins act both as adhesive molecules and as potential molecular effectors of the mechanical relay mechanism during wave propagation. These two roles are complementary. Indeed, adhesion to the vitelline membrane is essential to induce the early onset of tissue invagination and to sustain the 3D cycle of deformations. Moreover the mechanosensing properties of Integrins makes them candidates as molecular transducers to activate MyoII and transmit its activation from one cell to the next. It is remarkable that the spatially restricted expression of *scab* correlates with the specific mechanical sensitivity of endoderm cells (Ext.Fig.6).

The paradigm emerged over many years that genetic and biochemical patterns control cellular behaviours during morphogenesis^2^. However, the flow of information from biochemical inputs to force generation and deformation is typically considered to be unidirectional. Our observations reveal a striking departure from this paradigm, highlighting how spatial pattern and collective cell dynamics can emerge through tight dynamic coupling of mechanical feedback and biochemical activity. Another interesting example of this principle can be found in the zippering process in *Ciona intestinalis*^52^ whereby sequential MyoII contractions ahead of the zipper drive tissue deformations (new cell-cell contacts) that propagate signalling and contractility within a tissue. This opens the door to a deeper investigation of how mechano-chemical signalling not only induces cell shape changes but also gives rise to spatiotemporal patterns of deformation at the tissue level.

## Supporting information

Supplementary table 1 (genotypes and crosses)

## Author Contributions

A.B., C.C. and T.L conceived the project and planned experiments. A.B and C.C. performed all experiments and quantifications. A.B. did the simulations and discussed them with E.M. J-M.P. designed the MS2-fog construct and all other Sqh::mCherry. A.B., C.C. and T.L. analyzed the results and discussed them with E.M. P-F.L. gave inputs. A.B., C.C. and T.L. wrote the manuscript and all authors made comments.

## Acknowledgements

We thank all members of the Lecuit and Lenne group for stimulating and useful discussions during the course of this project. We thank the IBDM imaging facility for assistance with maintenance of the microscopes, FlyBase for maintaining curated databases and Bloomington for providing fly stocks. This work was supported by the ERC grant Biomecamorph #323027). We are grateful to Pavel Tomancak for sharing with us the information regarding the expression pattern of Scab published in [Ref 32] and its function in Tribolium. C.C. was supported by the CNRS, A.B. was supported by a PhD fellowship from the LabEx INFORM (ANR-11-LABX-0054) and of the A*MIDEX project (ANR-11-IDEX-0001–02), funded by the ‘Investissements d’Avenir French Government program’. E.M. was supported by NIGMS 1RO1 GM098441-06. We acknowledge the France-BioImaging infra-structure supported by the French National Research Agency (ANR–10–INBS-04-01, «Investments for the future»).

## Extended Figure legends

**Extended Figure 1.**
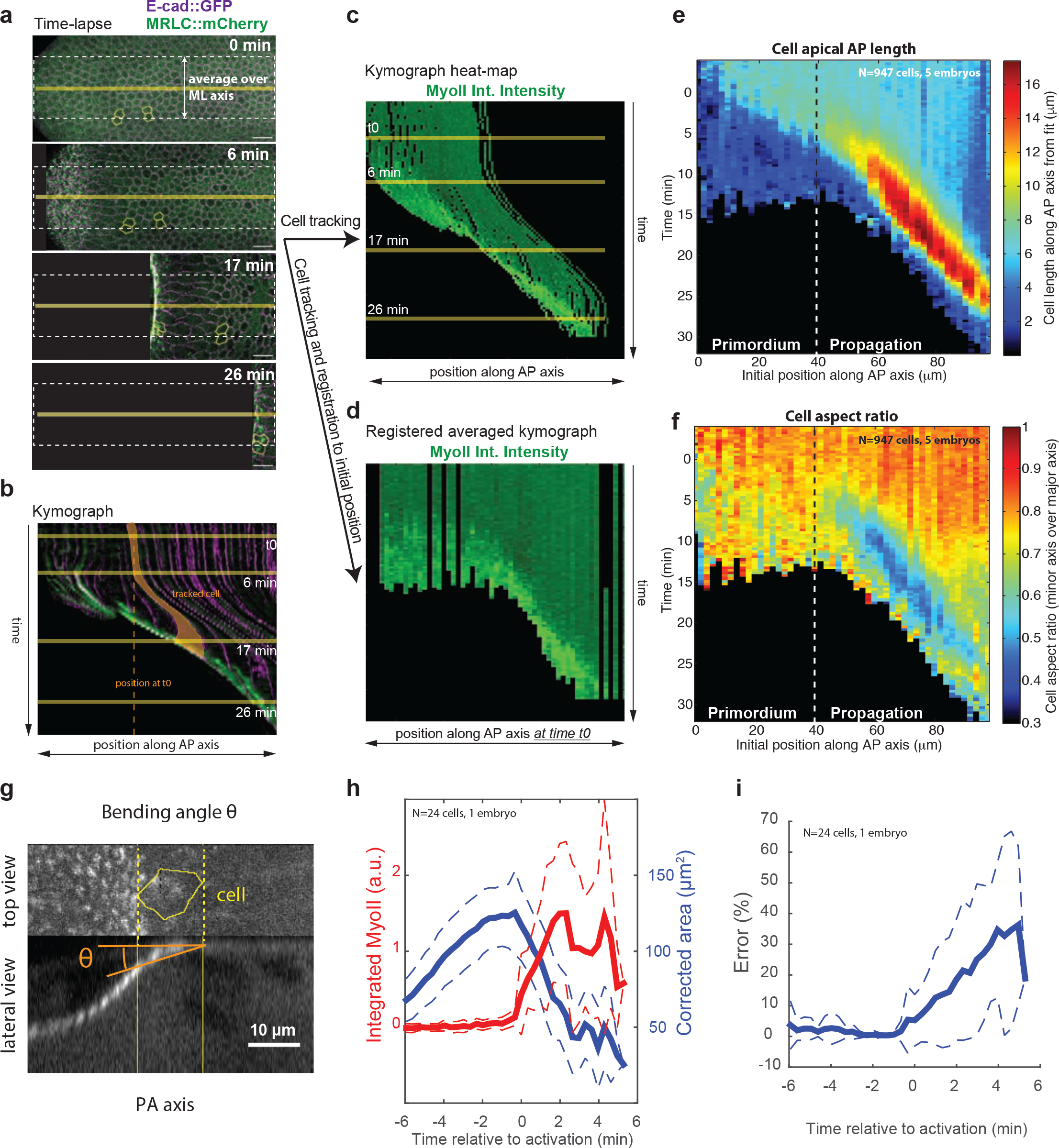
Quantifications of the morphogenetic wave propagation within the dorsal posterior epithelium. (**a-d**) Procedure to generate kymograph heat-maps from time-lapses of endoderm morphogenesis. In **a** stills of a dual color time-lapse, E-cad::GFP is used to label cell contours and MyoII is labeled with mCherry. Cell tracking is performed (few tracked cells are displayed in yellow) and cell positions and MyoII intensities are extracted over time. (**b**) A kymograph generated along the yellow horizontal line in **a**. A single cell, highlighted in yellow, first moves with the tissue towards the anterior and then increases its projected apical area before recruiting MyoII. Solid yellow lines indicate the time frames in **a** and the dashed vertical line the position of the cell at time t0. (**c**) Kymograph heat-map of MyoII integrated intensity extracted from cell tracking in **a**. Data are averaged along the ML (here shown for a single embryo) axis within the white dashed line box in **a** and cell positions are the positions at time t. Note that some visible individual cell tracks display movements similar to the kymograph in **b**. (**d**) Kymograph heat-map of MyoII integrated intensity registered on the cell positions at time t0 extracted from cell tracking in **a**. Data are averaged along the ML axis (here shown for a single embryo), as in **c**, but now each cell track is plotted according to its position along the AP axis at time t0. Note that some visible individual cell tracks do not display any movement and appear as straight vertical lines. As a result tissue deformation is not considered and the movement of the wave across the tissue can be visualized. This representation is used elsewhere in the manuscript. Time t0 is the onset of endoderm morphogenesis. (**e-f**) Kymograph heat-maps of AP apical cell projected length (**e**) and cell aspect ratio (**f**). The primordium and propagation regions are indicated. N=947 cells from 5 embryos. (**g**) Representation of the bending angle θ in a lateral view from a tracked cell (yellow contour) (see methods). (**h**) Average time traces of the corrected area (see methods) along with MyoII integrated intensity. Cells are registered on the time of the MyoII activation. Note that the corrected area decreases when MyoII intensity increases indicating that cells are truly constricting. (**i**) Time trace of the relative error in the measurement of cell area (i.e. underestimation of the real cell area) due to the bending angle. (**h-i**) Data from 24 cells, 1 embryo, with θ measured automatically (Method 1, see methods) on MyoII side views. Mean ± SD are shown.

**Extended Figure 2.**
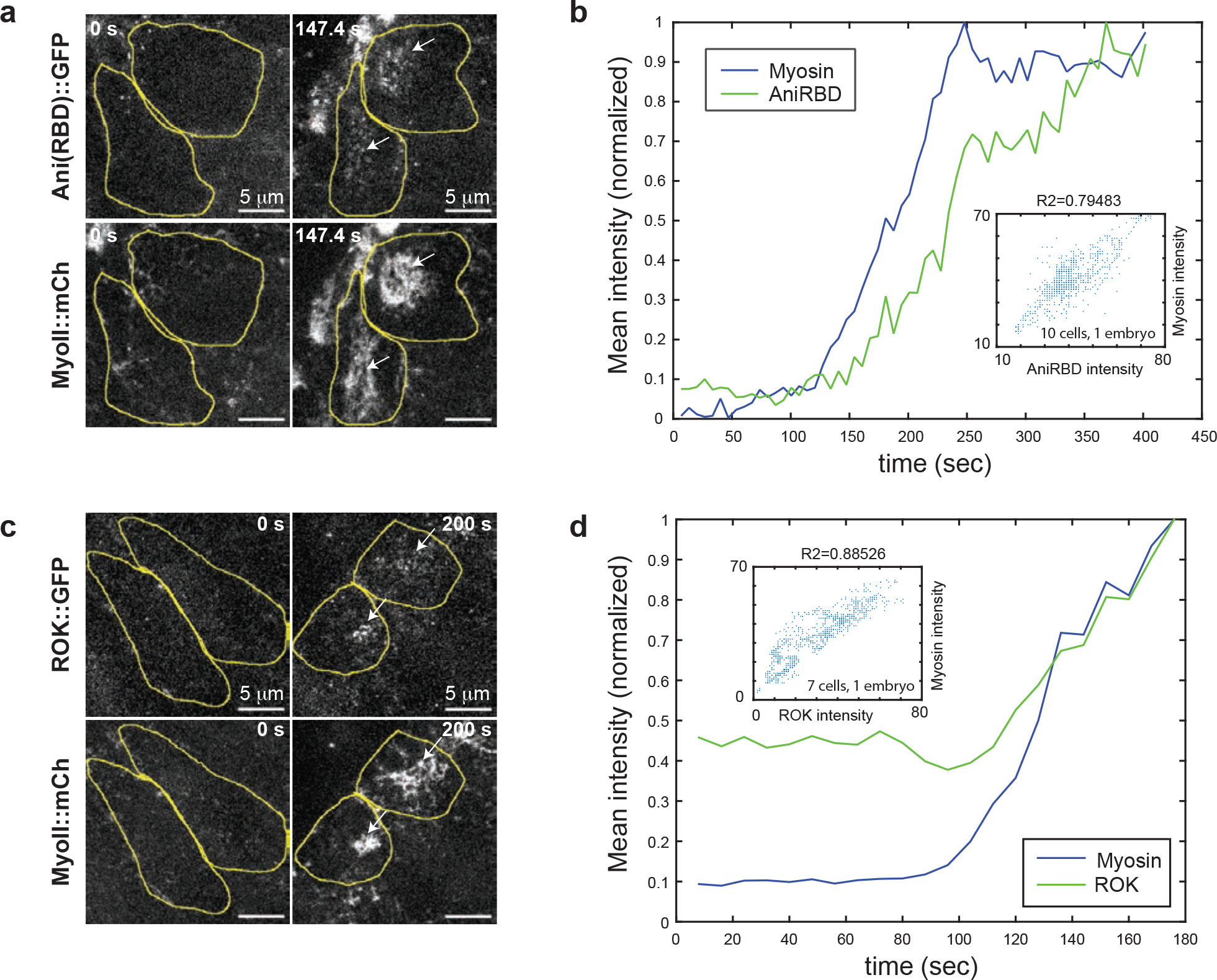
Rho-GTP and Rok propagate together with MyoII during the wave. (**a and c**) High resolution stills of cells in the propagation region labeled with MyoII and with the Rho1-GTP sensor, Anillin(RBD)::GFP (**a**), and RokKD::GFP (**c**) with the yellow line indicating cell contours. (**b and d**) Representative examples curves of Rho1 sensor (**b**) or MyoII and Rok (**d**) mean intensity over time in one cell of the propagation region. Intensity values are normalized to the max of each curve for visualization purposes. In the insets, scatter plots of MyoII intensity vs Rho1 sensor intensity (**b**) or Rok intensity (**d**) from ~10 cells of the embryo shown in (**a**) and ~7 cells of the embryo shown in (**c**) (see methods).

**Extended Figure 3.**
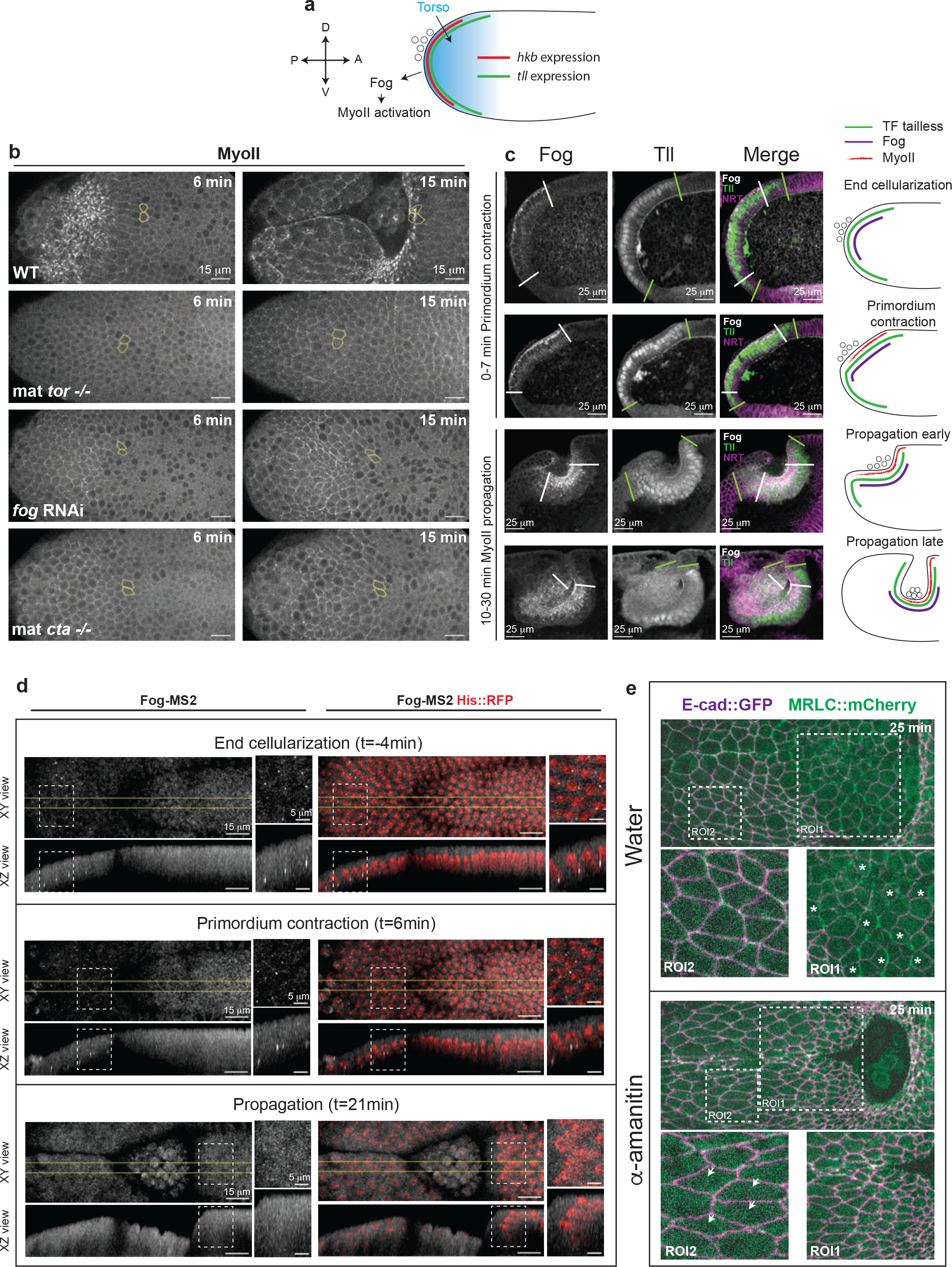
MyoII propagation is not controlled transcriptionally. (**a**) Schematics of the terminal patterning-dependent pathway controlling MyoII activation in the posterior endoderm. (**b**) MyoII activation in the posterior endoderm in embryos mutant for torso (mat *tor*−/−), depleted of maternal and zygotic *fog* (*fog* RNAi) and mutant for the Gα12/13 Concertina (mat *cta*−/−). Yellow contours mark cells in the propagation region. (**c**) Sagittal sections of embryos immunostained for Fog and Tll (together with the membrane marker Neurotactin in the merge) at different stages of posterior endoderm morphogenesis. On the right a cartoon representation of the distribution of the indicated molecules. (**d**) *fog* expression in the posterior endoderm visualized with the MS2-MCP system^35^ in living embryos. Top and side views of MCP::GFP and merge with Histone::RFP labelling nuclei are shown for the indicated time points. White dashed boxes indicate the positions of the close-ups on the right and the yellow lines the region where side-views were generated. (**e**) Stills of a control (water) and α-amanitin injected embryo at a late stage of invagination. Asterisks indicate dividing cells (ROI1) and arrowheads elongated cells in the posterior-ventrolateral ectoderm. Elongated cells in the ectoderm are a hallmark of cell stretching typical of conditions where cell intercalation is affected ^29,30,53^.

**Extended Figure 4.**
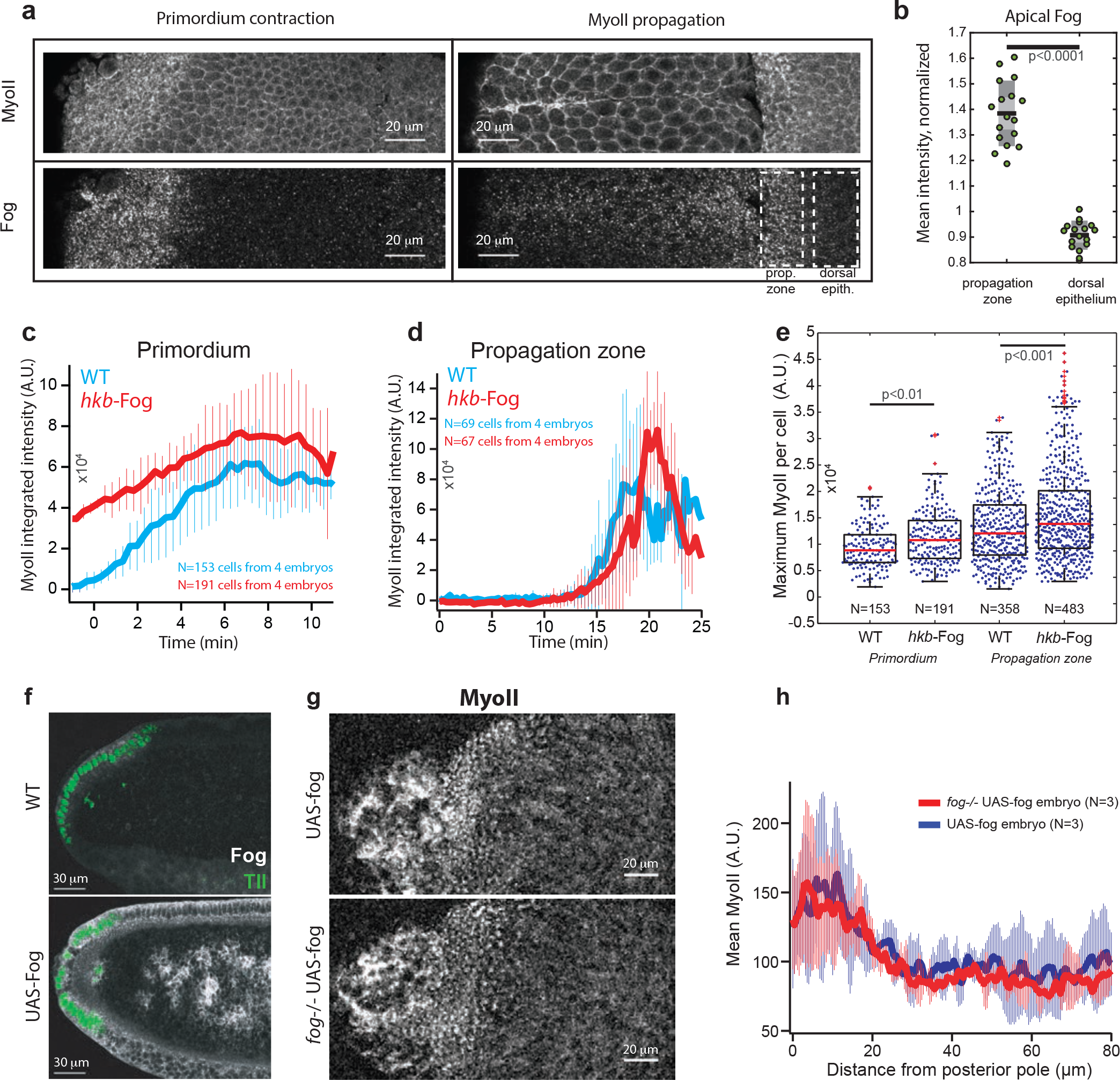
Fog diffusion and/or patterned Fog signaling do not control MyoII wave propagation. (**a**) Apical views of embryos stained for Fog and MyoII at primordium contraction and late propagation stages during endoderm morphogenesis. (**b**) Quantifications of Fog mean intensity in the propagation zone and in the dorsal epithelium. Individual embryos data points are superimposed to box plots (dark line: mean, light grey zone: 95% s.e.m., darker grey zone: s.d.). P<0.0001 from a Mann-Whitney test. (**c-d**) Time course of MyoII integrated intensity in cells of the primordium (**c**, N=153 and 191 cells for WT and *hkb-fog* from 4 embryos each) and in a 10 μm band of cells at ~30 μm distance from the primordium at time 0 (**d**, N=69 and 67 cells for WT and *hkb-fog* from 4 embryos each). Mean ± SD between different embryos are shown. (**e**) Maximum of MyoII integrated intensity in cells of the indicated conditions. Box plots are superimposed to individual data points. N indicates the number of cells from 4 embryos for each condition. P<0.01 and P<0.001 from a Mann-Whitney test. (**f**) Sagittal sections of a WT and an embryo ubiquitously expressing Fog (UAS-Fog). Immunolabelling for Fog is shown in white and for Tailless (Tll) in green. (**g**) MyoII pattern with Fog ubiquitous expression in a WT and a *fog* zygotic mutant embryos (imaged with 20X objective) (**h**) Quantifications of MyoII patterns along the AP-axis in the posterior endoderm of the indicated conditions. N=3 embryos each. In **c** and **d** time 0 min is the onset of endoderm morphogenesis (see methods).

**Extended Fig. 5.**
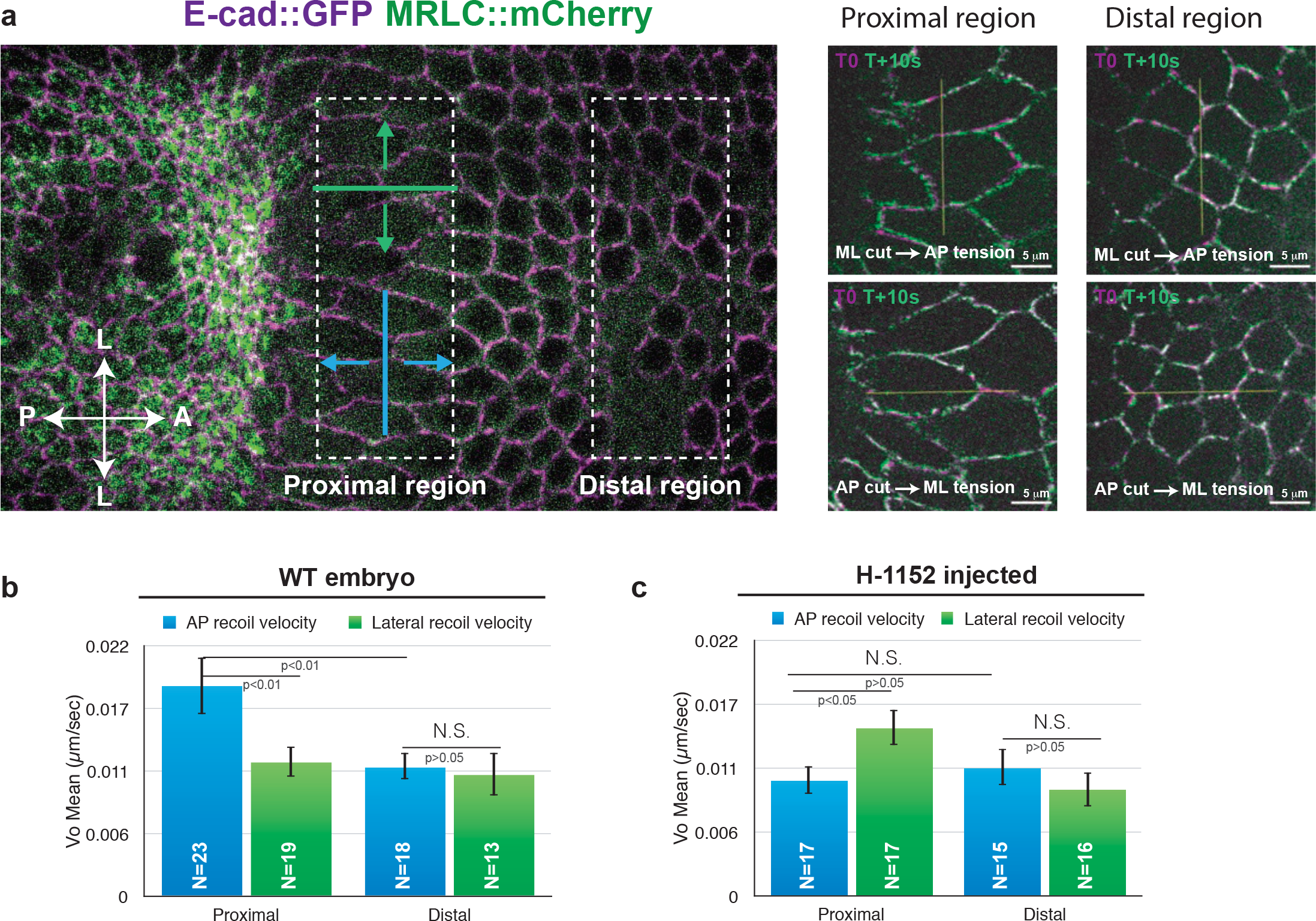
Cells in the propagation region are subjected to mechanical stress. (**a**) Left: Regions in the dorsal epithelium (corresponding to the propagation region) where tension was probed by line cuts with different orientations (ML cut in blue and AP cut in green). Right: Examples of ML and AP line cuts in the indicated regions. Overlays of the pre-cut (in magenta) and an image 10 s post-cut (in green) are shown. The yellow line indicates the line cut. (**b-c**) Quantifications of the tissue initial recoil velocity in the regions and line orientations illustrated in WT (**b**) and H-1152 (Rok inhibitor) injected embryos (**c**). N indicates the number of independent ablations extracted from 11 and 7 embryos for WT and H-1152 respectively. Mean values ± s.e.m. are shown.

**Extended Fig. 6.**
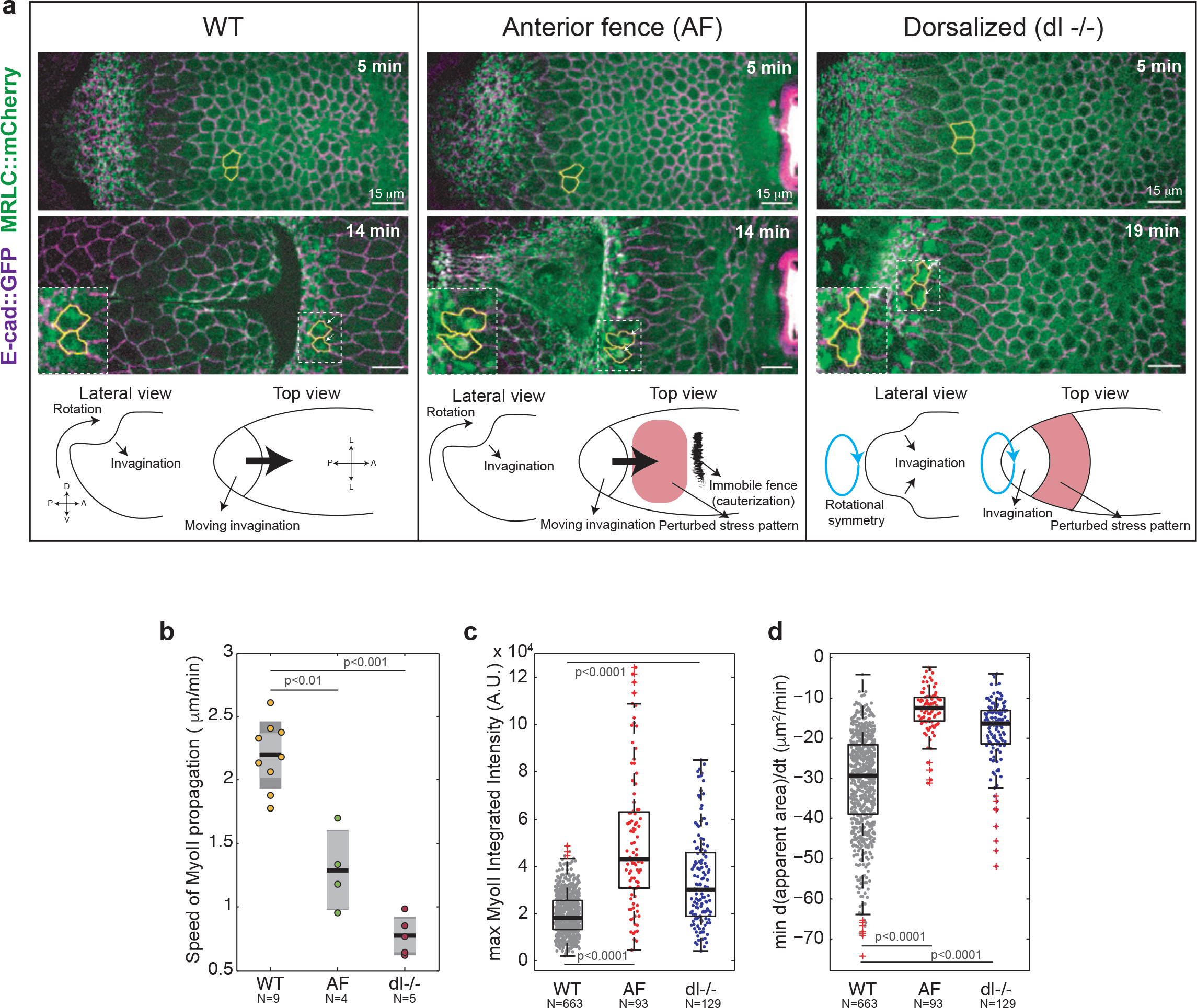
Mechanical perturbations affect propagation speed and MyoII recruitment. (**a**) Time-lapse stills of a WT (left), an embryo with an anterior medio-lateral fence (AF, middle) and a dorsalized embryo (dl−/−, right). MyoII::mCherry and cell contours labeled with Ecad::GFP are shown. Marked in yellow are representative tracked cells in the propagation region. Bottom: Schematic representations of the normal and perturbed conditions. (**b**) Speed of MyoII propagation in the reference frame of the tissue in the indicated conditions. Individual data points are superimposed to box plots (dark line: mean, light grey zone: 95% s.e.m., darker grey zone: s.d.). N=9 WT embryos, 4 AF embryos and 5 dl−/− embryos. P<0.01 and P<0.001 from a Mann-Whitney test. (**c-d**) Maximum MyoII integrated intensity (**c**) and minimum rate of apical constriction (**d**) in cells of the propagation region for the indicated conditions. Box plots superimposed to individual data points. N=663 cells from 9 WT embryos, 93 cells from 4 AF embryos and 129 cells from 5 dl−/− embryos. P<0.0001 from a Mann-Whitney test. In **a** time 0 is the onset of endoderm morphogenesis.

**Extended Fig. 7.**
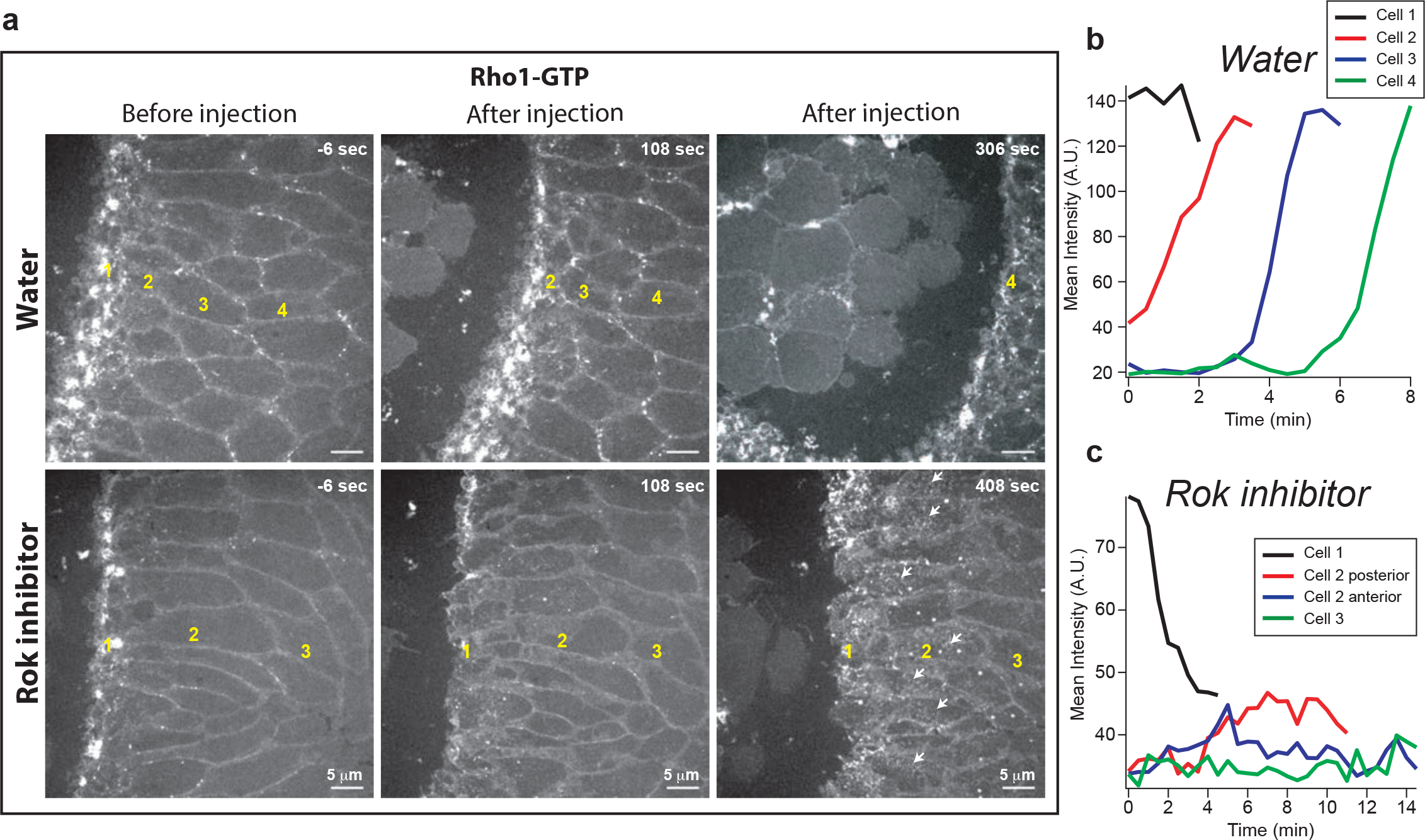
Rho1-GTP propagation is affected after Rok inhibitor injection. (**d**) Stills from a high resolution time lapse of Rho1 sensor (Anillin(RBD)::GFP) in a control (water) and an embryo injected with 40 mM H-1152 (Rok inhibitor) during propagation. Larger views and later times after injection are shown (compared to Fig.4c). The numbers in yellow indicate cells belonging to cell rows at different distances from the invaginating furrow. (**b-c**) Measurement of Rho1 sensor (Anillin(RBD)::GFP) mean intensity over time in representative examples of cells (or regions of cells) at different distances from the invaginating furrow in water (**b**) and H-1152 (**c**) injected embryos.

**Extended Fig. 8.**
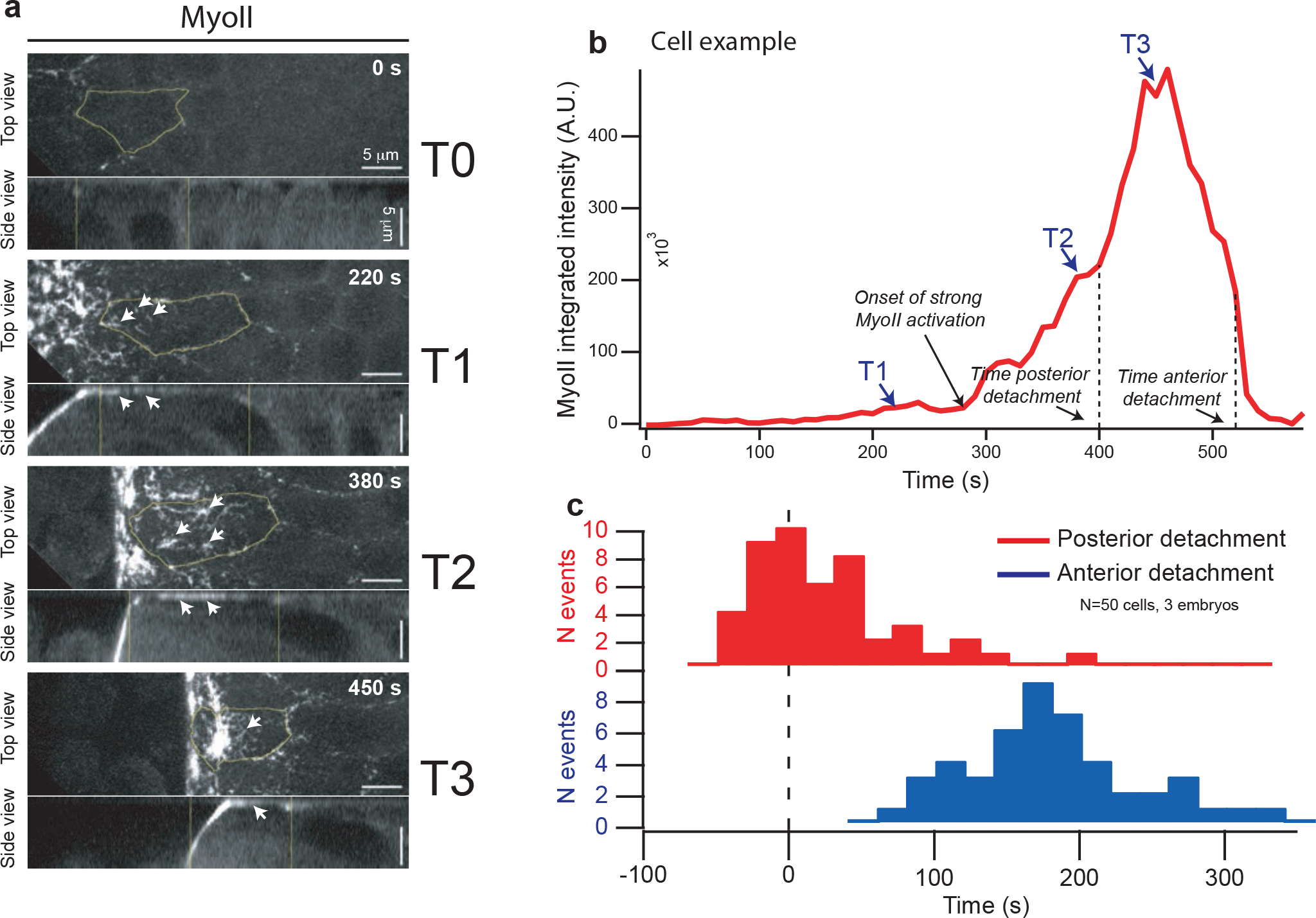
MyoII activation in cells of the propagation zone. (**a**) Stills from top and side views of MyoII activation in cells of the propagation region. The cell contour in labelled in yellow. White arrows indicate bright speckles of MyoII in regions of the cell in contact with the vitelline membrane. (**b**) Time trace of MyoII integrated intensity of the cell in **a**. The selected stills in **a** are indicated along with the time of cell posterior and anterior edge detachment. (**c**) Histogram of the time of cell posterior and anterior edge detachment from the vitelline membrane relatively to the time of MyoII activation (defined as time 0). N=50 cells from 3 embryos.

**Extended Fig. 9.**
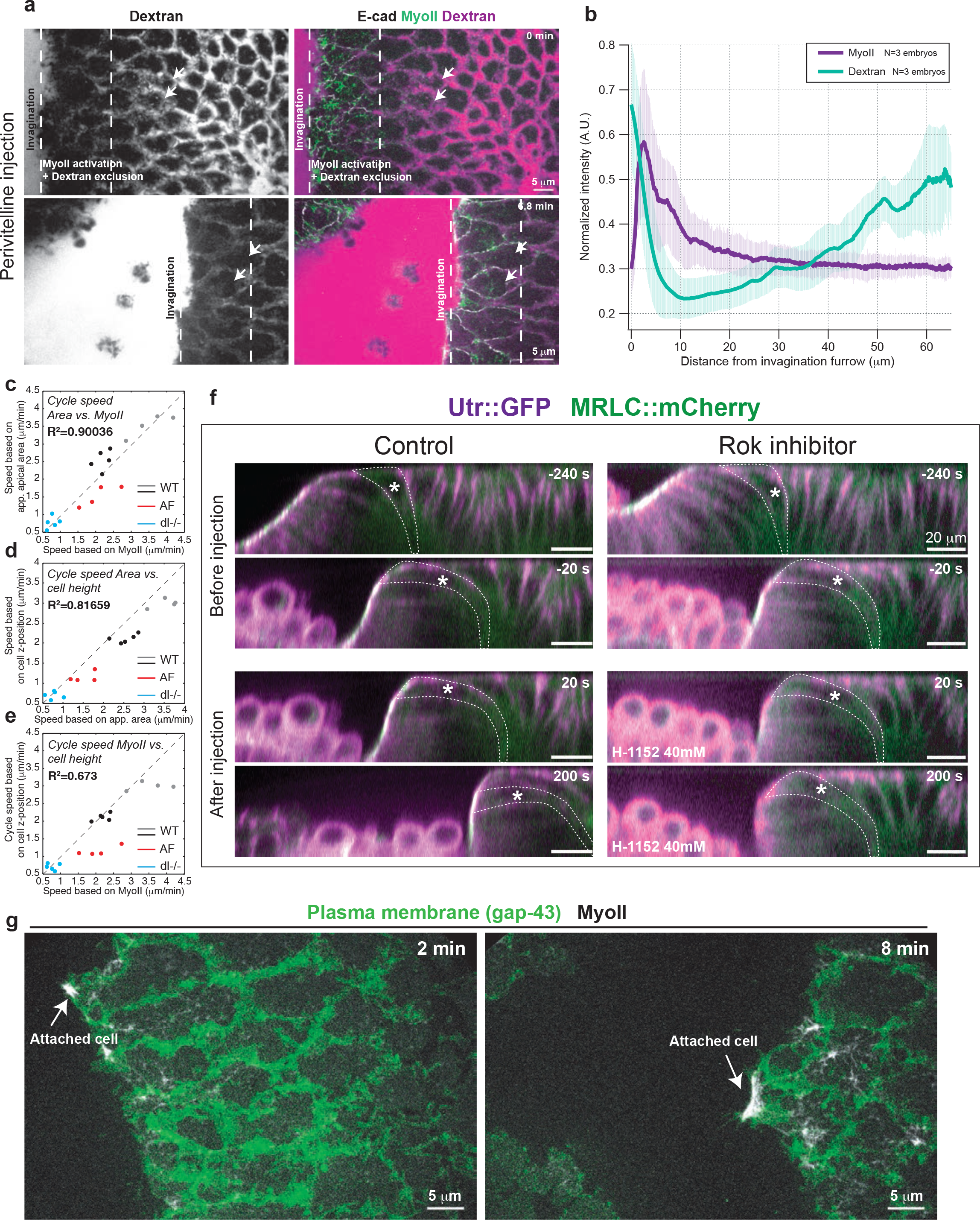
MyoII-dependent mechanical 3D cycle and cell attachment to the vitelline membrane in cells of the propagation zone. (**a**) Stills from a high-resolution time-lapse of dextran injection in the perivitelline space. Dextran alone is on the left and merge images with Ecad and MyoII on the right. Arrows indicate two cells as the invaginating furrows approaches. (**b**) Space-intensity plot of MyoII and dextran intensity in the propagation region. N=3 embryos. (**c-e**) Scatter plots of the average speed of the mechanical cycle during wave propagation calculated from the MyoII activation front vs the speed calculated from the projected apical cell area or cell position along the apico-basal axis of the tissue for the indicated conditions. R^2^ values for a fit y=x are shown. (**f**) A control and an embryo injected with H-1152 (Rok inhibitor) during MyoII propagation. Reconstructed side views from confocal stacks are shown. White asterisks and dashed lines mark a single cell over time. Time 0 is the time of injection. (**g**) Stills from a high-resolution time-lapse of MyoII and a plasma membrane marker (Gap-43) in cells of the dorsal epithelium during MyoII propagation. The white arrows indicate events of cell attachment to vitelline membrane.

## Movie Legends

**Movie 1**: Time-lapse of posterior endoderm morphogenesis in a WT embryo. E-cad::GFP is in magenta and MRLC::mCherry in green. The yellow contours mark tracked cells in the propagation zone. Scalebar 15 μm.

**Movie 2**: Time-lapse of MyoII recruitment in a cell in the primordium region (left) and in a cell in the propagation region, corresponding to cells in Fig. 1c and relative measurements in Fig. 1d. E-cad::GFP is in magenta and MRLC::mCherry in green. The cell contours are marked in yellow. Scalebar 3 μm.

**Movie 3**: High resolution time-lapse of cells in the propagation region labelled with the Rho1-GTP sensor Ani(RDB)::GFP (left) and MyoII (right). The white arrows point to Rho1-GTP and MyoII accumulations during wave propagation. Scalebar 5 μm.

**Movie 4**: High resolution time-lapse of cells in the propagation region labelled with Rok(KD)::GFP (left) and MyoII (right). The white arrows point to Rok and MyoII accumulations during wave propagation. Scalebar 5 μm.

**Movie 5**: Time-lapse of MyoII activation during posterior endoderm morphogenesis in a *torso* mutant embryo (laid by mothers *tor* −/−, top), in a strong knock-down of fog (fog RNAi, middle) and in a *concertina* (Gα12/13) mutant embryo (laid by mothers *cta* −/−, bottom). MyoII is labelled with MRLC::mCherry. Scalebar is 20 μm.

**Movie 6**: Top and side views of *fog* expression (MS2-*fog* revealed by MCP::GFP) on the left and MyoII (MRLC::mCherry) activation on the right during posterior endoderm morphogenesis in a WT embryo. Scalebar 15 μm.

**Movie 7**: Top and side views of *fog* expression (MS2-*fog* revealed by MCP::GFP) on the left and MyoII (MRLC::mCherry) activation on the right during posterior endoderm morphogenesis in a water (top) and α-amanitin (bottom) injected embryos. Scalebar 15 μm.

**Movie 8**: Time-lapse of MyoII activation and propagation during posterior endoderm morphogenesis in a water (top) and α-amanitin (bottom) injected embryos. E-cad::GFP is magenta and MRLC::mCherry in green. The yellow contours mark tracked cells in the propagation zone and the white arrows indicate cells at the moment of MyoII activation. Scalebar 20 μm.

**Movie 9**: Time-lapse of MyoII activation and propagation during posterior endoderm morphogenesis in a WT (top) and an embryo over-expressing *fog* in the primordium (hkb-Fog, bottom) injected embryos. E-cad::GFP is magenta and MRLC::mCherry in green. Time registration has been performed as described in the methods. Scalebar 20 μm.

**Movie 10**: Time-lapse of MyoII activation during posterior endoderm morphogenesis in embryos expressing ubiquitously *fog* (UAS-Fog, top), a constitutively active version of Gα12/13, (UAS-Cta[CA], middle) or a constitutively active version of Rho1 (UAS-Rho1[CA], bottom). MyoII is labelled with MRLC::mCherry. Scalebar 20 μm.

**Movie 11**: Rho1 and MyoII activation and propagation during posterior endoderm morphogenesis in a water injected embryo at end of cellularization. Rho1-GFP is detected with the Rho1 sensor Ani(RDB)::GFP (top) and MyoII is labelled with MRLC::mCherry (bottom). The white arrows indicate events of Rho1 and MyoII activation in cells of the propagation region. Scalebar 15 μm.

**Movie 12**: Rho1 and MyoII activation and propagation during posterior endoderm morphogenesis in an embryo injected with the Rok inhibitor H-1152 at the end of cellularization. Rho1-GFP is detected with the Rho1 sensor Ani(RDB)::GFP (top) and MyoII is labelled with MRLC::mCherry (bottom). The white arrows indicate events of Rho1 and MyoII activation in cells of the propagation region. Scalebar 15 μm.

**Movie 13**: Posterior endoderm morphogenesis in a WT embryo (top), an embryo with an anterior dorsal fence (middle) and a dorsalized embryo laid by *dorsal* −/− mothers (bottom). E-cad::GFP is in magenta and MRLC::mCherry in green. The yellow contours mark tracked cells in the propagation zone. Scalebar 15 μm.

**Movie 14**: Gastrulation in a WT (top) and a dorsalized embryo laid by *dorsal* −/− mothers (bottom). Scalebar 50 μm.

**Movie 15**: Time-lapse of a water (left) or Rok inhibitor (H-1152) injection during Rho1/MyoII wave propagation. Rho1-GFP is detected with the Rho1 sensor Ani(RDB)::GFP (top) and MyoII is labelled with MRLC::mCherry (bottom). Larger views and longer time lapse after injection. Note the slow and low-level activation of Rho1 in cells anterior to the invaginating furrow following Rok inhibition (top right). Scalebar 5 μm.

**Movie 16**: Time-lapse of a water (left) or Rok inhibitor (H-1152) injection during Rho1/MyoII wave propagation. Rho1-GFP is detected with the Rho1 sensor Ani(RDB)::GFP (top) and MyoII is labelled with MRLC::mCherry (bottom). Close-up on cells in the process of activating Rho1/MyoII at the time of injection. The white arrows point to Rho1-GTP and MyoII accumulations just before injection. The labels ‘Water’ and ‘H-1152’ mark the time of injection. Scalebar 5 μm.

**Movie 17**: Side view of MyoII recruitment and cell deformations during posterior endoderm morphogenesis. MyoII is labelled MRLC::GFP and the white arrows indicate events of MyoII activation during wave propagation. Scalebar 20 μm.

**Movie 18**: High-resolution time-lapse of MyoII (MRLC::mCherry) recruitment in a cell in the propagation region. Top and side views are shown. The cell contours are marked in yellow. At the bottom, a graph of MyoII integrated intensity with a moving vertical bar indicating the time in the time lapse. The white arrows point to bright accumulations of MyoII and the yellow arrows indicate the time of cell detachment at the posterior. Scalebar 5 μm.

**Movie 19**: High-resolution time-lapse of E-cad (E-cad::GFP) during apical cortex spreading onto the vitelline membrane in a cell in the propagation region. Top and side views are shown. Scalebar 5 μm.

**Movie 20**: Time lapse of a perivitelline injection of fluorescent Dextran labelling the extracellular space between cells and the vitelline membrane during MyoII wave propagation. Dextran is magenta, MRLC::mCherry is labelled in green and E-cad::GFP is in white. Scalebar 5 μm.

**Movie 21**: Time-lapse of a Rok inhibitor (H-1152) injection during MyoII propagation. Top and side views are shown. F-actin, labelling cell contours, is detected with Utrophin(ADB)::GFP (top) and MyoII is labelled with MRLC::mCherry (bottom). The label H-1152 marks the time of injection. Scalebar 15 μm.

**Movie 22**: Time-lapse of cells in the propagation region illustrating events of transient cell adherence to the vitelline membrane (seen as cells lagging behind the moving invaginating furrow and then suddenly snapping down in the invagination). MyoII, labelled by MRLC::GFP, is in white and cell plasma membrane, labelled by Gap-43::mCherry, is in green. Scalebar 5 μm.

**Movie 23**: Top and side views of posterior endoderm morphogenesis in a control (water injection) embryo (top), and an embryo injected with dsRNAs against *scab* (bottom). E-cad::GFP is in magenta and MRLC::mCherry in green. The yellow contours mark tracked cells in the propagation zone. Scalebar 15 μm.

## Materials and Methods

### Fly strains and genetics

The following mutant alleles and insertions were used: *tor*^*4*^, *dl*^*1*^, *fog*^*4a6*^, *cta*^*RC10*^, *sqh*^*AX3*^, *pUAST-fog*^*616*^, *pUAST-ctaQ303L* ^54^ *UAS-rho1V14* (Bloomington stock #8144), *hkb-fog* (chromosome 3 from^40^). MyoII RLC (MRLC) is encoded by the gene *spaghetti-squash* (*sqh*, Genebank ID: AY122159). Live cell imaging of Sqh was carried out using *sqh-Sqh::mCherry* (on chromosome 2 from ^5^ and a new insertions at the VK18 site, located 53B2 on chromosome 2 or the VK27 site located 89E11 on chromosome 3), and *sqh-Sqh::GFP* transgenics (gift from R. Karess). *E-Cad::EGFP* is a EGFP knock-in allele of E-Cadherin at the locus generated by homologous recombination (*E-cad::EGFP*^*Kin*^, ^55^). F-Actin was visualized using a *sqh-Utrophin::EGFP* insertion on chromosome 3^17^. Live imaging of the Rho1 sensor and Rok were carried out using a *ubi-Anilin(RBD)::EGFP* ^10^ and a *sqh-RokKD::GFP* ^9^(gift from Jennifer Zallen) transgenes. Plasma membrane was visualized using a *UASp-Gap43::mCherry* insertion (chr 3). *fog* gene expression was visualized with an MS2-*fog* pACMAN BAC inserted at the ATTp2 site, located 68A4 on chromosome 3 and the MCP-NoNLS::EGFP from ^35^. MCP-NoNLS::EGFP was provided maternally from females *;sqh-Sqh::mCherry;MCP-NoNLS::EGFP*, (insertion at VK18 site on chr 2) or *yw;Histone-RFP;MCP-NoNLS::EGFP* (Bloomington stock #60340) crossed with males carrying the MS2-*fog* BAC and F1 eggs were analyzed. *nos-Gal4* and *67-Gal4* (*mat α4-GAL-VP16*) are ubiquitous, maternally supplied.

For homogeneous expression of Fog, Cta[CA] and Rho1[CA] heterozygotes mothers *;67-Gal4,E-cad::EGFP*^*Kin*^,*sqh-Sqh:: Cherry;/+* were crossed with males carrying the *pUAST-fog*^6^, or *pUAST-ctaQ303L* or *UAS-rho1V14* transgene and F1 eggs were analyzed. The presence of the UAS-transgene was estimated on the basis of the phenotype. F1 progeny of males *Y/FM7 twi-Gal4, UAS-GFP;;UAS-fog^6^/+* crossed with female *fog*^*4a*^*/FM7 twi-Gal4 UAS-GFP;67-Gal4, E-cad::GFP^KIn^, sqh-Sqh::mCherry/+* was analyzed in Extended Fig. 4g-h. Mutant embryos were identified based on the absence of ubiquitous EGFP expression after imaging.

All fly constructs and genetics are listed in Supplementary Table 1.

### Constructs and transgenesis

<u>MS2-Fog construction:</u> the *fog* gene was modified in its 3’-UTR region (283 bp downstream the stop codon) by inserting 16 MS2 repeats (gift from Thomas Gregor^35^). A pACMAN BAC clone (CH321-09M06) which encompasses the 30kb of the *fog* gene along with 47kb upstream and 28kb downstream regions was modified by recombineering^56^. The recombined BAC was inserted in the genome at the attp40 landing sites (Chr2, 25C6), using PhiC31 mediated site-specific insertion transgenesis (BestGene, Inc.).

### RNA interference and drug injections

The dsRNA probe directed against *fog* was the one used in^57^. The dsRNA probe against *scab*, was 450bp long, and it is located in the coding region, targeting nucleotides 2652-3101 of scab-RC (Genebank ID: BT021944). dsRNA probes were prepared and injected at a final concentration of 5 μM in embryos less than 1 h old as previously described^29,57^.

α-Amanitin (from Sigma-Aldrich) was injected at a concentration of 500 mg/ml in embryos at the end of cellularization around 5 min prior imaging. The Rok inhibitor H-1152 (Enzo Lifesciences) was injected at a concentration of 40 mM in embryos either at the end of cellularization around 5 min before imaging or at stage 7 during imaging using an InjectMan4 micromanipulator and a FemtoJet 4i microinjector from Eppendorf directly installed on the imaging microscope.

Dextran injections were performed in the perivitelline space using Dextran 647 Alexa Fluor, 10000 MW, Anionic, Fixable from Thermo Fisher Scientific at a concentration of 1 mg/ml. Embryos were injected at stage 6 or 7 directly on the imaging microscope and imaged after few minutes (~2min) to allow diffusion of Dextran in the perivitelline space.

All injections were performed at 50-80% embryo length from the posterior and on the lateral side in all experiments in this paper.

### Live Imaging

For live imaging embryos were prepared as described earlier^58^ and imaged at stage 6 or 7 in the dorsal and posterior region for 10 to 60 min at room temperature (22°C) depending on the experiment. Most time-lapse imaging was performed with a dual camera spinning disc (CSU-X1, Yokogawa) Nikon Eclipse Ti inverted microscope (distributed by Roper) using a 20X/N.A. 0.75 water immersion, 40X/N.A. 1.25 water immersion or a 100X/N.A 1.4 oil-immersion objective depending on the experiment (20X or 40X magnification for tissue-level resolution and 100X magnification to obtain cellular level resolution). Dual color imaging of EGFP and mCherry/mKate2 FPs was performed by simultaneously exciting the fluorophores with a 491 nm and a 561 nm laser, and using a dichroic mirror to collect emission signals on two cameras.

Live imaging of laser ablation and laser cauterization experiments was performed with an Eclipse TE 2000-E microscope equipped with a spinning disc Ultraview ERS (Perkin-Elmer) using a 40×/N.A. 1.25 water-immersion, or a 100×/N.A. 1.4 oil-immersion objective. Dual color imaging of EGFP and mCherry was performed sequentially using a single camera by rapidly switching the excitation lasers (491 nm and a 561 nm) and emission filters.

For live imaging with 40X magnification z-series of 33-40 planes (spacing 0.8 μm) spanning ~30μm from the cell apex were acquired with a frame rate of 20s or 30s per frame. For live imaging with 100X magnification, z-series of 2-12 planes spanning 1 to 6 μm were acquired with a frame rate of 1-15s/frame. For laser ablations a single plane was imaged at a frame rate 0.5s/frame. Laser powers were measured at every imaging session and kept constant during the experiment.

Two-photon imaging of mid-sagittal sections (Fig. 5a) was performed with a Nikon confocal equipped with a Nikon 25X/N.A. 1.1 objective. A single section was acquired at a frame rate of 3s/frame.

### Generation of tissue fences by laser cauterization, and laser ablations

Tissue fences were generated within the dorsal epithelium along the Medio-Lateral (ML) embryonic axis at ~30% Embryo Length (EL) from the posterior as previously described^29^. Briefly, tissue cauterization and stitching to the vitelline membrane was obtained by focusing a near-infrared (NIR, 1030nm) laser with a 60X/N.A. 1.2 water-immersion Plan Apo VC, Nikon objective on the apical side of the embryonic blastoderm (~1–2 μm above adherens junctions) with an average power of 150 mW at the back aperture of the objective. To generate fences the stage was moved at a constant speed of 15–17 μm/s.

Line ablations were performed on the dorsal epithelium of embryos at stage 7 as previously described^29^. 20 μm line ablations were generated by moving the laser beam along the AP or ML axis within a region proximal (5–30 μm from the furrow) or more distal (60-100 μm from the furrow) to the invaginating furrow.

### Antibody staining

Embryos were fixed with 4% formaldehyde for 20 min, devitellinezed by shaking in Heptane/Methanol, and then stained according standard procedures^59^. Fog was detected with a Rabbit antibody anti-Fog (1:1000; gifted by Naoyuki Fuse^60^), Neurotactin with a mouse monoclonal antibody (1:100; BP-106 supernatant from DSHB), Tll with an anti-Tll polyclonal guinea pig antibody (1:250; ^61^, gifted by Cedric Maurange) and a chicken anti-GFP antibody was used to detect Sqh::EGFP (1:1000, AVES-GFP 120). The anti-Tll antibody was pre-adsorbed by incubation ON at 4C with fixed embryos less than 1h old. Secondary antibodies conjugates with Alexa488, Alexa555 and Alexa647 (from MoBiTec) were used 1:500. Stained embryos were mounted in Aqua-Polymount (Polysciences) and imaged with a Zeiss 510 confocal microscope using a C-Apochromat 40X/NA 1.2 or a 60X/NA 1.4 objectives. Image stacks with spacing of 0.3-0.5 μm were collected and sum projections of 3-7 planes analyzed.

### Image processing, segmentation and cell tracking

E-cad::GFP was used to label outlines and the apical side of the cells. A custom ImageJ macro integrating the Stack Focuser plugin from M. Umorin was used to project (by maximum intensity projection) a limited number of z-planes around the apical plane of cells (focused maximum projections). This resulted in sharper cell outlines and better S/N ratio compared with maximal projections. Briefly the macro uses the Stack Focuser plugin with a kernel size of 30 pixels (~ 6 μm) to identify the apical plane locally in all regions of the image on the basis of the E-cad::GFP signal. Then, only 4 planes for E-cad and 7 planes for MyoII around the identified apical plane are projected. The 2D projected stacks were then segmented and cells tracked as previously described^29^ using Ilastik (v1.2) and Tissue Analyzer^62^.

### Data analysis

All measurements of fluorescence intensity and cell shape parameters were extracted using the Fiji software and further analyzed with Matlab (including Curve Fitting Toolbox, Image Processing Toolbox, Statistics and Machine Learning Toolbox). To plot data the Igor software or Matlab (extended with custom designed functions and the UnivarScatter from Manuel Lera Ramírez) were used.

#### Measurements of fluorescence intensity in individual cells

Fluorescence intensities were measured on focused maximum intensity projections (for 40X image stacks) or standard maximum intensity projections (for 100X image stacks).

For individual cells measurements mean and integrated intensities were measured within an ROI obtained by automated segmentation and tracking and then shrunken by 4 or 10 pixels for 40X or 100X images respectively to remove junctional signals. For all images background intensities were subtracted by masking out relevant structures and then subtracting residual intensities. To generate the masks images were first background subtracted using a “rolling ball” algorithm (with a radius of ~6 μm for 40X images and ~1.2 μm for 100X images) and then an intensity threshold was used. Residual background images were smoothened with a Gaussian blur filter with a radius of 0.6-1 μm before subtraction.

Time registration of individual cells relative to MyoII activation was performed using a threshold on MyoII mean intensity.

The rates of MyoII recruitment in Fig.1i were calculated with a linear fit of the MyoII integrated intensity time traces within a window of 3 minutes after MyoII activation.

To extract maximum or minimum values of the integrated intensity (Ext.Fig.4e) and contraction rate (Ext.Fig.6d), the measurements of MyoII integrated intensity and projected area were smoothened over time with a rolling average using a window of 60sec.

For quantifications of Rho-GTP in Fig. 4f-g Ani(RBD)::GFP mean intensity was measured manually in circular ROIs of diameter 3.6μm. The rate of Rho1 sensor recruitment was estimated by a linear fit of mean intensity measurements in an interval of 60s before and 60s after injection.

#### Measurements of fluorescence intensity in the tissue

For the scatter plots in Extended Fig.2b and 2d, we measured mean MyoII and mean ROK or Rho sensor intensity in small regions of the tissue and plot one signal against the other. To do so, 100x movies of MyoII and ROK or Rho sensor where cropped respectively to 36×36μm^2^ or 25×25μm^2^ and registered using the Fiji plugin StackReg. Intensity was then averaged in bins of 3.6×3.6μm^2^ or 2.5×2.5μm^2^. Measurements of MyoII intensity vs ROK or Rho sensor intensity where plotted for all bins and all time frames (spanning ~5min) and correlation coefficient was calculated. We checked that binning did not affect the correlation.

For tissue level MyoII polarity measurements (Fig.3g), we used focused maximum intensity projections of embryos 10min after primordium activation. We subtracted background using a ‘rolling ball’ algorithm with radius 10μm and a threshold. We then measured mean MyoII intensity in rectangular ROIs in the active zone I(az) (which correspond to the propagation zone for WT and to the primordium for the other conditions) and the inactive zone intensity I(iz) (corresponding to the amniosierosa). The size of the ROIs spanned from ~10×30 μm^2^ to ~50×80 μm^2^ according to the visible area of the tissue (in WT the primordium has already invaginated at 10min). We defined polarity to be (I(az)-I(iz))/I(az).

For the space-intensity plot (Ext.Fig.9b), we measure MyoII and dextran mean intensity along ML axis and plot it as a function of the distance from the invaginating furrow along AP axis. Mean intensity of MyoII is measured after background was subtracted using a “rolling ball” algorithm (with radius 4μm). For both MyoII and dextran, measurements are smoothed along the AP axis with a rolling average using a window of 2μm. The position of the furrow was determined using the position where the derivative of the MyoII measurement first exceed its half-maximum. For each embryo data are averaged across several time points spanning 7-10min.

#### Measurement of cell bending angle

In order to estimate measurement artifacts due to the fact that the surface of cells is not parallel to the viewing plane, we measured the average angle between the cell surface and the imaging plane (or bending angle θ) for all time points. For this, we devised two methods (both using Matlab):

- <u>Method 1:</u> we reconstruct lateral views of the MyoII signal of tracked cells (see Ext. Fig.1g) and threshold them with a mean filter followed by erode/dilate binary operations. On the binary images of the cell profile, we obtain the coordinates of the cell surface by finding the maximum height of the cell *h* for each pixel position *x* along the AP axis. We then perform a linear fit *h*=*ax*+*b* and use it to calculate θ = tan^−1^(a) (defined positive for cells bending towards posterior).
- <u>Method 2:</u> we use the height information of the local apical plane in all regions of the image based of the E-cad::GFP signal (see methods for stack projections). This “height map” directly provides the coordinates of the cell surface that we average along the medio-lateral axis for each cell and on which we perform a fit similarly to the previous method.

The first method provides a reliable measurement of θ for the few cells from which we reconstructed lateral views (24 cells), while the second can be extensively applied to our dataset but underestimate the large bending angles due to space averaging performed to reconstruct the “height maps”. However, the two methods give values correlated with a Pearson coefficient of 0.55 and measure the same average error within 3 min after activation. Method 1 was used to measure θ in Extended Figure 1.

From θ, we correct the measurement of the cell surface using: true area=measured area/cos θ (see Ext. Fig.1h). We also estimate the relative error (i.e. the underestimation of the projected area compared to the real area) using error=100*(true area-measured area)/true area (see Ext. Fig.1i). Since the error is relatively small 2 minutes after activation (~20%), we decided to keep our projected area measurement as such. Note that we also measured the angle along the ML axis, but bending is very small along this axis (average max angle ~15° and error <3%).

#### Reconstruction of initial cell position for kymograph heat maps

In order to look at the propagation of cell shape and MyoII intensity parameters across the tissue, we elaborated kymograph heat maps to represent in a single graph space (position along the AP axis), time and value of any given parameter (area, integrated intensity, etc.). Measurements from cell tracking are used. Since at first approximation the phenomenon is symmetric along ML axis, we average data along this axis (Ext. Fig.1a and 1c). To remove the impact of cell movements (and estimate actual propagation from one cell to the next) we use initial coordinates for coordinates in the space axis (Registered kymograph Ext. Fig.1d). To do so, we bin individual cells in 2μm-wide bins according to their initial position along AP axis (their center position at time *t0* as found from tracking). If a cell *x* is not present in the first frame and appears only on frame number *tx*, we infer its initial position by extrapolating from the initial positions of all other neighboring cells present at *tx*. For any bin the median of the given parameter across all cells of the bin is used for color-coding. This representation can be used on any parameter measured from cell: integrated intensity, area, velocity, etc and enables to monitor propagation within the tissue independently of tissue deformation.

#### Measurements of the wave speed

We measured the wave speed either automatically from tracked cells data, or, when segmentation and tracking were not performed, by manual counting.

##### Automatic measurement

Speed of the wave can be measured either in the reference frame of the tissue (or initial frame reference, i.e. time t0) as defined for kymograph heat maps, or in the reference frame of the microscope. In Ext.Fig.6b, we compare the speed of the wave across the tissue (from one cell to the next) in conditions where tissue deformations are very different, hence we use the first method. In Fig.3d, we compare the speed of the wave set by extracellular diffusion in conditions where deformations are very similar, hence we use the second method.

To measure the speed of the MyoII activation wave in the reference frame of the tissue (initial frame reference), we first binarize the kymograph heat maps of the mean intensity of MyoII, select manually the region corresponding to the propagation (spanning around 20min corresponding to 10-30 min from the onset of endoderm morphogenesis) and then perform a linear fit of the positions of MyoII activation as a function of their time of activation. Activation is defined by mean MyoII intensity crossing a threshold. We proceed similarly for measuring the speed of the mechanical cycle using cell apical area and cell position along the apico-basal axis of the tissue (position in z), we use the reconstructed kymograph heat maps displaying cell apical area, or the position in z of the cadherin belt as a proxy for the cell surface (Ext.Fig.9c-e).

To measure the speed of the MyoII activation wave in the reference frame of the microscope, we perform a linear fit of the cell position at the time of MyoII activation as a function of time of activation. Here the positions are read directly from the microscopy images. Activation is defined by mean MyoII intensity crossing a threshold. Such a reference frame is used in Fig.3d where we are interested in diffusion of Fog in the perivitelline fluid.

##### Manual measurement

To obtain the wave speed in the reference frame of the tissue (initial frame of reference), we manually recorded the time of MyoII activation cells of the propagation zone belonging to 3-4 rows of cells aligned along AP-axis in each embryo. For each embryo we averaged over the different rows, and different embryos were averaged to obtain plots of the number of cells having activated MyoII along an AP row as a function of time. Time 0 is defined as the time of activation of the first cell in the propagation zone. This measurement is used in Fig. 2d, and using the Rho1-GTP sensor instead of MyoII, in Fig. 4b. We also extract the number of cells having activated MyoII within 20min from these measurements (Fig.3h).

#### Measurements of tension

The average tissue tension in Extended Fig. 5, proportional to the initial outward velocity after line cuts, was measured as previously described^29^ by particle image velocimetry (PIV).

### Simulations of wave propagation by diffusion or mechanical feedback

Full details on the simulations can be found in the Supplementary Methods.

### Embryo synchronization

Embryos synchronization was performed to compare different mutant conditions with WT embryos and to register embryos in developmental time before pooling and averaging. Synchronization is based on two developmentally-regulated events: 1) the disappearance of MyoII from the cellularization front at the basal side of cells and 2) the onset of cell divisions in the dorsal posterior ectoderm (see Ext. Fig. 3e). Both events are independent of morphogenesis of the posterior endoderm. Time t0 is then defined as the onset of MyoII recruitment and apical constriction in WT embryos.

For immunofluorescence experiments in fixed samples embryos staging was determined manually on the basis of similarity with events observed in time-lapse experiments.

### Statistics

For all experiments data points from different cells/embryos from at least 2 independent experiments were pooled to estimate the mean, s.d. and s.e.m. values plotted. All of the *P* values are calculated using a two sided non-parametric Mann–Whitney test (Matlab statistic toolbox) except in Fig.3g where P values are calculated using a one-sample t-test (null hypothesis of a normal distribution with mean equal to zero, i.e no tissue-level polarity of MyoII, Matlab statistic toolbox). No statistical method was used to predetermine sample size. The experiments were not randomized, and the investigators were not blinded to allocation during experiments and outcome assessment.

### Repeatability

All measurements were performed in 1–11 embryos. In experiments where embryos are not injected we consider each embryo as an independent experiment. In drug and dsRNA injection experiments the number of independent experiments is defined as the number of independent injections. Representative images, which are shown in Figs 1–6 and Extended Figs 1–9 were repeated at least twice and up to more than ten times.

### Data Availability

The authors declare that the data supporting the findings of this study are available within the paper and its supplementary information files. Raw image data are available upon reasonable request.

### Code availability

The custom codes used to process images analyze data and run simulations are available upon request.

