## Supplementary Information for "Transcriptional induction and mechanical propagation of a morphogenetic wave"

### Supplementary Methods

#### Simulations of the Myosin-II wave

In the following, we introduce simple toy models and simulations to describe and compare two general classes of mechanism that could generate a wave of Myosin-II (MyoII) activation throughout a tissue. For simplicity, we use one-dimensional (1D) models and evaluate the minimal ingredients necessary to reproduce the essential features of the MyoII wave, namely a sharp wavefront of MyoII concentration travelling at constant speed.

The first model we describe is based solely on chemical reaction-diffusion: a ligand (Fog) diffuses and binds irreversibly to its receptor, and receptor-bound Fog activates MyoII. In this model, ligand diffusion and local saturation of receptor binding are minimal ingredients required to reproduce the observed wave of MyoII. In the second model we describe the tissue as an active viscous medium, which can deform and flow in response to forces produced by MyoII. In this model, MyoII is locally activated within a source region by the Fog ligand, but the requirements for wave propagation are force transmission through the medium, and positive feedback via stress-dependent activation of MyoII.

### 1 Models

#### 1.1 Reaction-diffusion model (Related to Fig.3)

A previous reaction-diffusion model [1] obtained steep travelling waves of receptor-bound ligand concentration. Thus we first consider a similar simple 1D model in which Fog concentration is governed by local production, diffusion, and receptor binding, and receptor-bound Fog acts locally to activate MyoII. We introduce the following variables:

**Concentration of free Fog ligand $f$:** We assume that Fog is secreted only within a source domain $S$ (defined by its position $-x_p \leq x \leq x_p$), at a rate $k_s$. We assume further that Fog diffuses away from this source with diffusion coefficient $D$.

**Concentration of receptor-bound Fog $f_b$:** We assume that free Fog ligands bind irreversibly at a rate $k_b$ to a receptor that is present at an initially homogeneous concentration $R^0$. We assume that the total amount of receptor is constant (on the time scale of the contractile wave). Therefore the concentration of receptors that are available for binding is $R^0 - f_b$.

**Active MyoII concentration** $m$**:** We assume that MyoII is activated at a rate proportional to receptor-bound Fog $f_b$, with rate constant $k_a$, and that it is inactivated at a rate proportional to its concentration with rate constant $k_i$.

Given these assumptions, the time evolution of free Fog concentration $f$, bound Fog $f_b$ and MyoII $m$ is governed by the system of equations:

$$\begin{cases} \partial_t f = D\partial_x^2 f + k_s S - k_b(R^0 - f_b)f \\ \partial_t f_b = k_b(R^0 - f_b)f \\ \partial_t m = k_a f_b - k_i m \end{cases} \tag{1}$$

With

$$S(x) = \begin{cases} 1 & -x_p \leq x \leq x_p \\ 0 & \text{otherwise} \end{cases} \tag{2}$$

##### 1.2 Mechanochemical model (Related to Fig.4)

Next, we consider a mechanochemical model in which MyoII is present within a material which has the ability to deform or flow. Fog ligand activates MyoII locally within a source domain, and MyoII activity spreads through transmission of MyoII-dependent stress through the material and positive feedback via stress-dependent activation of MyoII.

###### Mechanics of the tissue

Previous theoretical work has detailed different ways in which contraction waves could propagate within a multicellular tissue [2] or the actomyosin cortex [3][4]. Here, we develop a minimal model of contraction waves in a simplified tissue. We consider a 1D active viscous fluid with effective viscosity $\eta$, with active stress $\sigma$ generated proportionally to local MyoII concentration $m$, and whose motions are resisted by friction with an external medium. We note that a viscoelastic version of this model yields similar results.

Following [5], we write the constitutive equation for an active viscous fluid as:

$$\sigma = Cm + \eta\partial_x v \tag{3}$$

Where $m$ is local activity of MyoII and $C$ a constant. We assume that the only external force is dynamic friction, proportional to the velocity of the material $v$ with friction coefficient $\gamma$. Neglecting inertial forces, forces balance instantaneously as:

$$\partial_x \sigma - \gamma v = 0 \tag{4}$$

The velocity field of the tissue is then given by:

$$\partial_x^2 v - \frac{1}{l^2} v = -k_m \partial_x m \tag{5}$$

Where $l = \sqrt{\eta/\gamma}$ is the hydrodynamic length, and $k_m = C/\eta$.

The time evolution of velocity $v$ is thus set by the time evolution of $m$.

##### MyoII activation

We replace the previous model for ligand diffusion, receptor binding and MyoII activation with a simpler model of an activating signal $f$, which is produced only within the source domain $S(t)$ and advected (reflecting displacement of the primordium with tissue deformation), and locally activates MyoII via a saturating Hill function of the signal $f$ with Hill coefficient $n$. In addition, we assume a simple form of mechanical feedback in which MyoII is activated by local stress $\sigma$ also via a saturating Hill function. Finally, we assume that MyoII is advected by flow of the tissue (with velocity $v$), and inactivated at a constant rate $k_i$. Accordingly, the equations (1) for $f$ and $m$ are written as:

$$\begin{cases} \partial_t f = k_s S - \partial_x (vf) \\ \partial_t m = k_a \frac{f^n}{f^n + 1} + k_f \frac{\sigma^n}{\sigma^n + \theta_f} - k_i m - \partial_x (vm) \end{cases} \tag{6}$$

Note that the boundary of the primordium (domain $S$ where the signal $f$ is produced) is now moving with the contraction of the tissue such that its position $x_p$ evolves with time (see eq.2). Besides, the signal $f$ does not diffuse. Thus the sole role of $f$ is to initiate MyoII activation locally, while the spread of MyoII activity beyond the primordium must be governed by stress transmission and stress-dependent activation of MyoII.

### 2 Simulations

#### 2.1 Discretization

We used Eulerian coordinates (i.e. fixed positions) for space coordinates.

To integrate equations (6) numerically, we used a central difference approximation in space (where the error is proportional to the chosen discretization space step $\Delta x^2$) and either an Euler (where the accumulated error is proportional to the chosen time step $\Delta t$, for the mechanochemical model) or a Runge-Kutta 4 (to $\Delta t^4$, for the reaction-diffusion model) time integration method.

To solve equations (5), we computed analytically its Green function, as the solution of:

$$\partial_x^2 v - \frac{1}{l^2} v = \delta_0 \tag{7}$$

with boundary conditions $v(-x_{max}) = 0$ and $v(x_{max}) = 0$. To obtain the velocity field at each time, we then convolved this Green function $G$ with $-k_m \partial_x m$ (the right-hand side of eq (5)):

$$v(x,t) = -k_m \int_{-\infty}^{\infty} G(x-u)\partial_x m(u,t) du \tag{8}$$

We used central differences for the calculation of the spatial derivative.

#### 2.2 Implementation and parameters

We used Matlab for implementation, including its built-in function for convolution.

##### Units

Units of concentration were fixed either by the initial concentration of receptor $R^0$ (in the chemical model) or by the ratio between activation and inactivation rates $k_a/k_i$ (in the mechanochemical model). Concerning length and time, we fixed the maximum simulated time to be $t_{max} = 20min$ and the primordium border to be at $x_p = 40\mu m$, to be commensurate with the experiments.

##### Chemical parameters

In the reaction-diffusion model, local saturation of the receptor is required for a sharp wave front to propagate [1]. To satisfy this requirement, receptor binding must be sufficiently fast relative to diffusion to achieve efficient saturation. Note that a simpler model in which Fog would directly activate MyoII (without the intermediate of the receptor-bound Fog) can only spread MyoII activity with a wave front that flattens away from the source.

We chose rates $k_b$, $k_s$ and $k_a$ to reproduce a full activation of MyoII in the primordium in $\sim 5min$, and we chose $D = 20\mu m^2.sec^{-1}$ to obtain a wave speed of $\sim 7\mu m.min^{-1}$ as measured in experiments.

In the mechanochemical model, the propagation of contraction waves requires non linearity (a Hill function coefficient $n = 2$ is sufficient), sufficiently strong positive feedback ($k_f$), and a threshold of stress ($\theta_f$) small enough to obtain saturation of the Hill function in the range of stress accessible by the system. When these minimal conditions are satisfied, the wave speed is set by feedback strength and activation threshold - higher feedback strength and lower threshold values yield faster waves. Accordingly, we chose $k_f$ and $\theta_f$ to reproduce the observed wavespeed of $\sim 7\mu m.min^{-1}$.

##### Mechanical parameters

We set the hydrodynamic length to $l = 20\mu m$, representing the distance from the MyoII front to the position at which deformation occurs. We chose $k_m$ sufficiently small to prevent collapse of the tissue within the simulation time but big enough to obtain a primordium contraction $> 50\%$.

#### 2.3 Comparison with experiments

To compare the results of the simulations with the results of our experiments, we generated similar plots as the ones done for experimental measurements (heat-map kymographs, see methods). Note that we aimed here for a qualitative, rather than a quantitative, comparison.

##### Position and speed of activation front with time

We defined activation time $t_a(x)$ for a given position $x$ as the first time point at which MyoII concentration $m(x,t)$ exceeds the value 0.5. We then plotted the position of activation $x$ as a function of time of activation $t_a$ (see Fig.3b).

To obtain the speed, we fitted the positions of activation as a function of time by a second degree polynomial $x(t_a) = at_a^2 + bt_a + c$ and took its derivative. Since $a$ happened to be non zero, the speed varied with time. However, we could still compare the overall speed in different conditions.

##### Registered kymograph heat-maps

When no deformation is present, kymograph heat-maps can be produced by simply displaying $m(x,t)$. This is not the case otherwise. Our simulations use Eulerian coordinates (time and fixed positions). As such, they are well suited for viscous flows, but they are not the most suited to keep track of large deformations with time (as would naturally be Lagrangian coordinates). To compare with the experimental kymograph heat-maps, we added the stored deformation field $e(x,t)$ [7] to our simulation in order to record deformation of the 1D material (or tissue). It is evolving with time following:

$$\partial_t e = \partial_x v + e\partial_x v - v\partial_x e \tag{9}$$

By integrating this deformation field in space for each point $x$, we obtained initial positions $x_0(x,t)$ and thus we could generate kymograph heat-maps in initial reference frame, as for experiments. More precisely, we used:

$$x_0(x,t) = \int_{u=0}^{u=x(t)} \frac{1}{1+e(u,t)} du \tag{10}$$

**MyoII integrated intensity**

In the kymograph heat-maps obtained from experiments, we displayed the MyoII signal integrated over one cell. However, in our simulations we computed the MyoII concentration $m(x,t)$. To better compare both kymograph heat-maps, we displayed for simulations the concentration of MyoII integrated over the unit volume taking into account its local deformation.

We thus calculated the local amount of MyoII $M(x,t)$ for kymograph heat-maps (Fig.4e) as follows:

$$M = m\Delta x(e+1) \tag{11}$$

Where $\Delta x$ is the space discretization step and $e(x,t)$ the stored deformation .
